# Shared and unique lifetime stressor characteristics and network connectivity predict adolescent anxiety and depression

**DOI:** 10.1101/2024.10.25.620373

**Authors:** Yueyue Lydia Qu, Sidhant Chopra, Shijie Qu, Carrisa V. Cocuzza, Loïc Labache, Clemens C.C. Bauer, Francesca Morfini, Susan Whitfield-Gabrieli, George M. Slavich, Jutta Joormann, Avram J. Holmes

## Abstract

**Background:** Exposure to major life stressors and aberrant brain functioning have been related to anxiety and depression, especially during adolescence. However, whether these associations differ based on the specific characteristics of the stressors experienced, and/or the functional networks engaged, remains unclear.

**Methods:** We used baseline lifetime stressor exposure and resting-state functional magnetic resonance imaging data from a longitudinal sample of 150 adolescents enriched for anxiety and depressive disorders. We examined the cumulative lifetime stressor frequency and severity of five stressor characteristics: physical danger, interpersonal loss, humiliation, entrapment, and role change/disruption. Anxiety and depression symptoms were assessed at three time points: baseline, 6-month and 12-month follow-ups. Linear mixed-effect models tested if the lifetime frequency and severity of these stressor characteristics and functional connectivity within and between frontoparietal, default, and ventral attention networks at baseline predicted anxiety and depression symptoms at three time points.

**Results:** Lifetime frequency and severity of humiliation and entrapment predicted both anxiety and depression symptoms. Lifetime frequency and severity of entrapment exposures predicted anxiety and depression symptoms after accounting for baseline depression and anxiety symptoms, respectively. Resting-state functional connectivity between default, frontoparietal and ventral attention networks did not predict either anxiety or depression symptoms after correcting for multiple comparisons.

**Conclusions:** Our study highlights lifetime exposures to humiliation and entrapment stressors as central stressor characteristics predictive of prospective anxiety and depression symptoms in adolescence. Our results also suggest that resting-state functional connectivity within and between default, frontoparietal and ventral attention networks may be relatively weak predictors of prospective anxiety and depression symptoms in adolescence.

## Introduction

Exposure to major life stressors is a strong risk factor for the onset and subsequent recurrence of affective disorders (Faravelli & Pallanti, 1989; Francis et al., 2012; Kendler et al., 1999; Kessler, 1997; Miloyan et al., 2018), especially in adolescence when there is increased brain plasticity and heightened vulnerability to the emergence of psychopathology (Casey et al., 2008; March-Llanes et al., 2017; Merikangas et al., 2010, 2010; Paus et al., 2008). However, stressors come in many different forms and are diverse with respect to both their characteristics and associations with mental health outcomes (Bonanno et al., 2023; Cohen et al., 2019; A. H. Lee et al., 2025; Slavich, 2019). Results from large-scale, prospective cohort studies have suggested that exposures to certain characteristics of major life stressors may preferentially increase risk for specific clinical outcomes such as anxiety and depression. For instance, stressor characteristics that involve devaluation of the self, such as interpersonal loss and humiliation, are theorized to preferentially heighten risk for depression (Asselmann et al., 2015; Farmer & McGuffin, 2003; Finlay-Jones & Brown, 1981; Keller et al., 2007; Kendler et al., 2003). In contrast, stressor characteristics marked by a threat to one’s physical integrity, such as danger, are theorized to be stronger predictors of anxiety (Asselmann et al., 2015; Ayazi et al., 2014; Finlay-Jones & Brown, 1981; Kendler et al., 2003). Stressors characterized by feelings of failure without any means of escape, such as entrapment, predict both anxiety and depression (Griffiths et al., 2014; Kendler et al., 2003). Although distinct stressor characteristics are not mutually exclusive from each other (e.g. a stressor can be both dangerous and humiliating), prior studies have often examined each stressor characteristic in isolation. Therefore, the *unique contributions* of each stressor characteristics to the prediction of anxiety and depression symptoms on top of each other remains unclear. Furthermore, very few studies examined the associations between lifetime stressor exposures and psychopathology in adolescence.

With respect to the brain, prior studies have linked altered brain functioning with anxiety and depression in early life. Specifically, resting-state functional connectivity (RSFC) within and between the frontoparietal, default and ventral attention networks, which are theorized to underlie dysfunctional executive (Cole et al., 2014; Kaiser et al., 2015; Menon, 2011; Williams & Goldstein-Piekarski, 2020; Xu et al., 2019; Z. Zhang et al., 2025), self-referential (Berman et al., 2011; Kaiser et al., 2015; Menon, 2011; Williams & Goldstein-Piekarski, 2020; Xu et al., 2019; Z. Zhang et al., 2025) and salience processing (Manoliu et al., 2014; Menon, 2011; Williams & Goldstein-Piekarski, 2020; Xu et al., 2019) respectively, have been associated with both anxiety and depression in pre-adolescent children and adolescents (Ho et al., 2015; Pawlak et al., 2022; Perino et al., 2021; Rzepa & McCabe, 2018; Sievertsen et al., 2025; Willinger et al., 2024).

However, only a few studies have examined RSFC correlates of anxiety and depression symptoms using longitudinal samples of pre-adolescent children and adolescents (Connolly et al., 2017; Huang et al., 2023; Jalbrzikowski et al., 2017; Jin et al., 2020; Lopez et al., 2018; Pawlak et al., 2022; Qu et al., 2021; Strikwerda-Brown et al., 2014). Therefore, it remains unclear if certain RSFC patterns may precede changes in anxiety and depression symptoms during these critical developmental periods.

### Present Study

To address these gaps in knowledge, we first investigated if lifetime exposures (self-reported number of exposures [henceforth referred as “frequency”] and subjective severity ratings [henceforth referred as “severity”]) to any characteristics of major life stressors assessed at baseline predicted anxiety and depression symptoms at each of the three time points— baseline, 6-month and 12-month follow-up assessments. We then analyzed if any RSFC metric within and between the frontoparietal, default, and ventral attention networks assessed at baseline predicted anxiety and depression symptoms at these three time points. These analyses were conducted using linear mixed-effects (LME) models in a longitudinal sample of adolescents recruited from school-based and hospital-based child treatment programs (Hubbard et al., 2024). Most participants had a current diagnosis of at least one anxiety or depressive disorder at the time of baseline assessment. Our results showed that lifetime frequency and severity of humiliation and entrapment stressors predicted both anxiety and depression symptoms over time. Furthermore, lifetime frequency and severity of entrapment exposures are differentially associated with prospective anxiety and depression symptoms, respectively. These findings indicate that among all stressor characteristics we assessed, humiliation and entrapment exposures are the strongest predictors of anxiety and depression in adolescence.

Whether such exposures are differentially associated with specific mental health outcomes may depend on the exact measure (frequency or severity) used. Although our study showed that RSFC within the default network, between default and frontoparietal networks and between ventral attention and frontoparietal networks predicted prospective anxiety symptoms, the main effects of these associations did not survive the false discovery rate (FDR) corrections. These findings indicate that RSFC within and between the default, frontoparietal, and ventral attention networks may have limited predictability for prospective anxiety and depression symptoms in adolescence.

## Methods and Materials

### Participants

Data were collected from 215 adolescents (*M*_age_ = 15.44, range = 14-17 years old) enrolled in the Boston Adolescent Neuroimaging of Depression and Anxiety (BANDA) study (Hubbard et al., 2024) and assessed at 6-month intervals after the initial visit for up to 12 months. The present analyses were restricted to data from the baseline, 6-month follow-up and 12-month follow-up assessments. Resting-state functional magnetic resonance imaging (rsfMRI) data were available for 202 participants at baseline (Hubbard et al., 2024). Out of these 202 adolescents, 150 had available behavioral measures of interest at baseline, 6- and 12-month follow-up assessments and were included in the final sample for statistical analyses involving stressor characteristics and symptom measures (**Table 1**). Of these 150 participants, 65% (*n* = 98) had a current diagnosis of at least one anxiety or depressive disorder. Diagnostic group of each participant was given by a blinded, licensed clinical psychologist based on the Diagnostic and Statistical Manual of Mental Health Disorders, 5^th^ Edition (DSM-5; American Psychiatric Association, 2013) and reached moderate to substantial inter-rater agreement (Hubbard et al., 2024).

**Table 1.** Demographic characteristics for the final analytical sample (*n* = 150)

| Baseline |  |  |
| --- | --- | --- |
| Demographics | *M/n* | *SD/%* |
| Age | 15.44 | 0.85 |
| Sex, (% female) | 92 | 61.33 |
| Mean parental education <sup>a</sup> | 6.11 | 1.27 |
| Diagnostic group | *n* | % |
| Anxiety <sup>b</sup> | 58 | 38.67 |
| Control | 52 | 34.67 |
| Depression <sup>c</sup> | 40 | 26.67 |
| Race | *n* | % |
| African American | 3 | 2.00 |
| Asian | 3 | 2.00 |
| Hawaiian | 1 | 0.67 |
| White | 121 | 80.67 |
| Ethnicity | *n* | % |
| Hispanic | 11 | 7.33 |
| Lifetime frequency of stressor characteristics | *M* | *SD* |
| Entrapment | 4.36 | 3.18 |
| (Range: 0-14; Possible range: 0-28) |  |  |
| Humiliation | 5.85 | 4.60 |
| (Range: 0-20; Possible range: 0-40) |  |  |
| Interpersonal loss | 5.08 | 3.92 |
| (Range: 0-20; Possible range: 0-61) |  |  |
| Physical danger | 1.98 | 2.18 |
| (Range: 0-10; Possible range: 0-46) |  |  |
| Role change/disruption | 2.99 | 2.47 |
| (Range: 0-15; Possible range: 0-24) |  |  |
| Lifetime severity ratings of stressor characteristics | *M* | *SD* |
| Entrapment | 13.65 | 12.01 |
| (Range: 0-54; Possible range: 0-95) |  |  |
| Humiliation | 8.88 | 7.92 |
| (Range: 0-36; Possible range: 0-60) |  |  |
| Interpersonal loss | 12.09 | 9.57 |
| (Range: 0-45; Possible range: 0-85) |  |  |
| Physical danger | 5.87 | 6.81 |
| (Range: 0-38; Possible range: 0-60) |  |  |
| Role change/disruption | 7.20 | 6.51 |
| (Range: 0-32; Possible range: 0-60) |  |  |
| Clinical symptoms | *M* | *SD* |
| Anxiety symptoms | 26.05 | 18.24 |
| (Range: 0-73; Possible range: 0-93) |  |  |
| Depression symptoms | 16.84 | 15.28 |
| (Range: 0-56; Possible range: 0-66) |  |  |
| Six-month follow-up |  |  |
| Clinical symptoms | *M* | *SD* |
| Anxiety symptoms | 24.86 | 17.51 |
| (Range: 0-79; Possible range: 0-93) |  |  |
| Depression symptoms | 17.14 | 14.60 |
| (Range: 0-58; Possible range: 0-66) |  |  |
| Twelve-month follow-up |  |  |
| Clinical symptoms | *M* | *SD* |
| Anxiety symptoms (0-93) | 21.72 | 15.24 |
| (Range: 0-62; Possible range: 0-93) |  |  |
| Depression symptoms (0-66) | 14.95 | 13.90 |
| (Range: 0-53; Possible range: 0-66) |  |  |

Note: <sup>a</sup> Mean parental education = the average of the highest education degree achieved across both parents for each subject. <sup>b</sup> Anxiety = having a current diagnosis of at least one anxiety disorder and no depressive disorder based on DSM-5. <sup>c</sup> Depression = having a current diagnosis of at least one depressive disorder based on DSM-5.

### Measures of social-psychological characteristics of life stressors

At the baseline assessment, the Stress and Adversity Inventory for Adolescents (STRAIN; Slavich et al., 2019) assessed each participant’s total lifetime frequency and severity ratings of exposures to five social-psychological stressor characteristics—physical danger, interpersonal loss, humiliation, entrapment, and role change/disruption—by quantifying both the frequency and subjective severity ratings of self-reported acute and chronic stressors across the life course (see https://www.strainsetup.com). Examples of stressors linked to each of these five characteristics have been described elsewhere (Slavich et al., 2019). The mean lifetime frequency and severity of each stressor characteristic within each diagnostic group at baseline assessment are illustrated in **Supplementary Figure 1**. Self-reported severity ratings is a good complementary measure to frequency of major life stressors because subjective ratings do not assume that all stressors are perceived equally by all individuals and may thus index important individual differences in stress vulnerability (Shields et al., 2023). Total lifetime frequency and severity for these five characteristics were used to predict the levels of anxiety and depression symptoms at each of the three time points (baseline, 6-month and 12-month follow-up assessments). Brief definitions for these stressor characteristics are as follows (Brown et al., 1995; Finlay-Jones & Brown, 1981; Kendler et al., 2003; Slotter & and Walsh, 2017):

### Physical danger

The degree of potential future threat to one’s physical safety that might occur as a result of the stressor.

### Interpersonal loss

Diminution of a sense of connectedness or well-being as a result of a real or realistically imagined loss of a person by death or by separation.

### Humiliation

The likelihood of a stressor rendering a person devalued in relation to others or self, usually due to rejection or a sense of core failure.

### Entrapment

Ongoing circumstances of marked difficulty of at least 6 months’ duration that the individual can reasonably expect to persist or get worse, with little or no possibility that a resolution can be achieved as a result of anything that might reasonably be done.

### Role change/disruption

Life transitions that involve addition, subtraction or change of social roles.

The STRAIN has demonstrated excellent test-retest reliability, good concurrent and discriminant validity, as well as predictive utility in relation to various clinical outcomes including anxiety and depression (Slavich et al., 2019; Slavich & Shields, 2018).

### Measures of anxiety and depression symptoms

For each participant, anxiety and depression symptoms were assessed at three time points: baseline, 6-month and 12-month follow-ups. The mean anxiety and depressive symptoms within each diagnostic group at each of the three time points is illustrated in **Supplementary Figure 2**.

Self-reported anxiety symptoms were assessed by the Revised Children’s Anxiety and Depression Scale (RCADS; Chorpita et al., 2000). The RCADS has exhibited excellent internal consistency, test-retest reliability, convergent and discriminant validity (Chorpita et al., 2000). The RCADS have six subscales: separation anxiety disorder, social phobia, generalized anxiety, panic disorder, obsessive-compulsive disorder and low mood. Total anxiety symptoms for each participant were computed by summing the four anxiety subscales (Separation Anxiety Disorder, Social Phobia, Generalized Anxiety and Panic Disorder).

Self-reported depression symptoms were assessed by the Mood and Feelings Questionnaire (MFQ; Angold et al., 1995; Costello & Angold, 1988). Prior studies have found the MFQ to be a reliable and valid measure of adolescent depression in both clinical and non-clinical samples across different populations (Burleson Daviss et al., 2006; Sund et al., 2001; Thabrew et al., 2018; Wood et al., 1995). We used the total MFQ score as the measure of depression symptoms for each participant.

### Neuroimaging

#### Data acquisition and processing

Functional and anatomical neuroimaging data were acquired at baseline assessment using a 3-Tesla Siemens Prisma scanner with a 2D multi-band gradient-recalled echo-planar imaging (EPI) sequence. Each participant underwent four 5.8-minute resting-state functional MRI (rsfMRI) runs, consisting of two runs with opposite phase encoding directions (AP/PA). Each rsfMRI scan was acquired using 2mm isotropic resolution and a TR of 800ms. Full details of the acquisition protocol can be found elsewhere (Siless et al., 2020).

The acquired rsfMRI data then went through the previously established Human Connectome Project (HCP) minimal preprocessing pipelines (Glasser et al., 2013). Minimally preprocessed T1w images (Glasser et al., 2013) went through bias- and distortion-correction using the *PreFreeSurfer* pipeline and registered to MNI space. Cortical surface reconstruction was conducted using FreeSurfer v5.2 using recon-all adapted for high-resolution images. The reconstructed surface meshes were then registered to the Conte69 surface template (Van Essen et al., 2012). During preprocessing, the fMRI data were first corrected for gradient-nonlinearity-induced distortions. The fMRI time series in each frame were then realigned to the single-band reference image to correct for subject motion using rigid body transformation (Jenkinson et al., 2002; Jenkinson & Smith, 2001) with FSL. The resulting single-band image underwent spline interpolation to correct for distortions and was then registered to the T1w image (Greve & Fischl, 2009). The registered fMRI volumes then went through nonlinear registration to the Conte69 surface template (Van Essen et al., 2012) and mapped to the standard CIFTI grayordinate coordinate space. Further details about the HCP minimal preprocessing pipelines of structural and functional images can be found elsewhere (Glasser et al., 2013). The minimally preprocessed rsfMRI data for each run were then denoised using ICA+FIX (Griffanti et al., 2014; Salimi-Khorshidi et al., 2014) pre-trained using HCP_hp2000.RData and aligned across participants using MSMAll multi-modal surface registration (Glasser et al., 2016; Robinson et al., 2014). The ICA-FIXed rsfMRI data were further processed by detrending and regressing out the 12 motion parameters within each run using customized Python code. Out of the 202 participants with available rsfMRI data, we excluded participants (*n* = 51) if their mean framewise displacement (Power et al., 2012) was greater than 0.30mm, if more than 20% of their volumes had FD > 0.30 mm, or if any single volume exceeded 5mm FD (Parkes et al., 2018; Satterthwaite et al., 2013). Consequently, our final sample for statistical analyses involving RSFC and symptom measures comprised 114 participants.

#### Resting-state functional connectivity

We defined 400 cortical regions of interest (ROIs) using the Schaefer’s Parcellation (Schaefer et al., 2018). Within each of the 4 rsfMRI runs, resting-state functional connectivity (RSFC) was measured by Pearson’s *r* correlations between the mean time series of each pair of ROIs. The average FC matrix across all runs in each participant was computed after applying Fisher Z-transformation and used for subsequent analyses.

Specifically, this study focused on RSFC within and between the frontoparietal, default, and ventral attention networks (**Figure 1**) according to the 17-network solution (Thomas Yeo et al., 2011) as predictors of anxiety and depression symptoms at three time points (baseline, 6-month follow-up and 12-month follow-up). Within-network connectivity was assessed by averaging the pairwise RSFC of all regions assigned to that network, resulting in three within-network connectivity values per individual. “Between” network connectivity was assessed by computing the pair-wise correlations of each ROI in one network (e.g., frontoparietal) to each ROI in the other network (e.g., default) and averaging across them, resulting in three between network connectivity values per individual.

**Figure 1.**
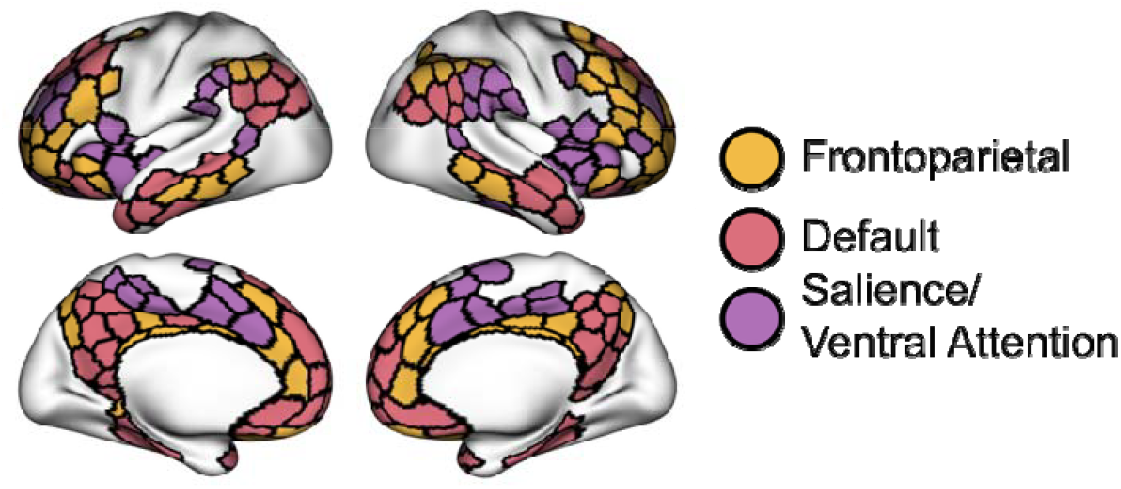
The functional network organization of the human cerebral cortex revealed through intrinsic functional connectivity. Colors reflect regions estimated to be within the same network. Cortical regions-of-interest (ROIs) defined by Schaefer’s parcellation (Schaefer et al., 2018) and assigned to frontoparietal (yellow), default (red), and salience/ventral attention (purple) networks (Thomas Yeo et al., 2011).

### Covariates

The following covariates were dummy coded, converted to factors and entered into each LME model: participant’s ethnicity (Hispanic: Yes=1, No=0), sex (female=0, male=1). The following covariates were continuous and entered into each LME model: The mean parental education (computed as the average of the highest education degrees across both parents), participant’s age at baseline. We further included baseline depression symptoms in each LME model predicting anxiety symptoms and baseline anxiety symptoms in each LME model predicting depression symptoms as continuous covariates to parse out differential predictors of each symptom (i.e., anxiety, depression) over time.

### Imputation of missing data

Missing values in the behavioral data were imputed using the *missForest* algorithm (Stekhoven & Bühlmann, 2012), an iterative imputation method based on random forest. The *missForest* algorithm was selected for its ability to impute both numeric and factor data and for its superior performance compared to 15 other common imputation algorithms as evaluated by the *missCompare* framework (Varga, 2020). Under the *missCompare* framework, 50 simulated datasets matching multivariate characteristics and missingness patterns of the original data were generated. The missing data in each simulated dataset were then imputed using a curated list of 16 algorithms. The computation time and imputation accuracy of each algorithm were hitherto assessed by calculating Root Mean Square Error (RMSE), Mean Absolute Error (MAE) and Kolmogorov-Smirnov (KS) values between the imputed and the simulated data points (**Supplementary Figure 3**). These metrics were compared under three conditions: Missing Completely At Random (MCAR), Missing At Random (MAR), and Missing Not At Random or Non-Ignorable (MNAR). Finally, post-imputation diagnostics, including visual examination of data distributions and stability of correlation coefficients between measures, before and after imputation, were examined, showing minimal impact of imputation (**Supplementary Figure 4**).

### Statistical analyses

We constructed linear mixed-effect (LME) models with random intercepts using the *lme4* package in Rv4.2.0 (Bates et al., 2015) to assess if lifetime frequency and severity of any stressor characteristics and functional connectivity within and between frontoparietal, default, and ventral attention networks at baseline assessment predicted each symptom (i.e., anxiety, depression) at each of the three time points (baseline, 6-month and 12-month follow-up assessments). Each LME model used one of the following three sets of predictors: (a) total lifetime frequency of five stressor characteristics [formula:

*Symptom* ~ *PhysicalDangerFrequency* + *InterpersonalLossFrequency* + *HumiliationFrequency + Entrapment Frequency + RoleDisruptionFrequency + BaselineAge + Sex + Ethnicity + MeanParentalEducation + Time + (*1|*SubjectID)];* (b) total lifetime severity ratings of five stressor characteristics [formula: *Symptom* ~ *PhysicalDangerSeverity + InterpersonalLossSeverity + Humiliationseverity + Entrapmen tSe verity + RoleDisruptionSeverity + BaselineAge + Sex + Ethnicity + MeanParentalEducation + Time +* (1|*SubjectID*)]; and (c) each of the six RSFC metrics within and between the frontoparietal, default, and ventral attention networks [formula: *Symptom*~*RS FC withinFPN (or RSFCwithinDN, RSFCwithinVAN, RSFCbetwFPN-DN, RSFCbetwFPN-VAN, RSFCbetwDN-VAN) + BaselineAge + Sex + Ethnicity + MeanParentalEducation + Time +*(1|*SubjectID*)] to predict anxiety or depression symptoms at each of the three time points, resulting in sixteen LME models in total. We accounted for covariates in each LME model to assess if the fixed effects of these stressor characteristics and RSFC metrics were robust to the inclusion of potential confounders. Continuous predictors and covariates were standardized to make the beta estimates more interpretable. We used the participant ID as the random intercept, meaning that the intercept estimates were allowed to vary across participants while the beta estimate for each predictor was constant across participants.

Despite the strong intercorrelations among the five stressor characteristics (*r*s > 0.50; **Supplemental Table 1**), LME models including all stressor characteristics as predictors showed variance inflation factors (VIFs) smaller than 5, demonstrating no multicollinearity issues. However, when the six RSFC metrics within and between the three functional networks were included as predictors in the same LME models, we observed moderate multicollinearity issues (4 < *VIF*s < 10). Hence, we entered these RSFC metric as separate predictors of anxiety and depression symptoms (*VIF*s < 2). Since the model residuals exhibit non-normality and heteroscedasticity (**Supplementary Figures 5-8**), square-root transformation was applied to the symptom variables to achieve normality and homoscedasticity (**Supplementary Figures 9-12**). Hence, all results presented and discussed in this manuscript are based on square-root transformations. For ease of interpretation, we back-transformed the beta coefficient associated with each bivariate association.

## Results

### Greater total lifetime frequency and severity of humiliation and entrapment exposures predict higher prospective anxiety and depression symptoms

We first used total lifetime frequency and severity ratings of all stressor social-psychological characteristics (i.e., physical danger, interpersonal loss, humiliation, entrapment, role change/disruption) at baseline to predict each symptom (i.e., anxiety, depression) at each of the three time points (baseline, 6-month and 12-month follow-up assessments). After accounting for lifetime frequency of four other stressor characteristics and other potential confounding covariates, greater total lifetime frequency of humiliation and entrapment exposures at baseline assessment prospectively predicted higher anxiety (Humiliation: *β* = 0.35, *β*_*back-transformed*_ = 0.18, *p*_*uncorrected*_ = 0.0073, *p*_*FDR*_ = 0.018; Entrapment: *β* = 0.78, *β*_*back-transformed*_ = 0.42, *p*_*uncorrected*_ = 2.87 × 10^−7^, *p*_*FDR*_ = 1.44 × 10^−6^, **Figure 2**) and depression (Humiliation: *β* = 0.38, *β*_*back-transformed*_ = 0.20, *p*_*uncorrected*_ = 0.0042, *p*_*FDR*_ = 0.011; Entrapment: *β* = 0.77, *β*_*back-transformed*_ = 0.39, *p*_*uncorrected*_ = 1.15 × 10^−6^, *p*_*FDR*_ = 5.73 × 10^−6^, **Figure 2**) symptoms.

**Figure 2.**
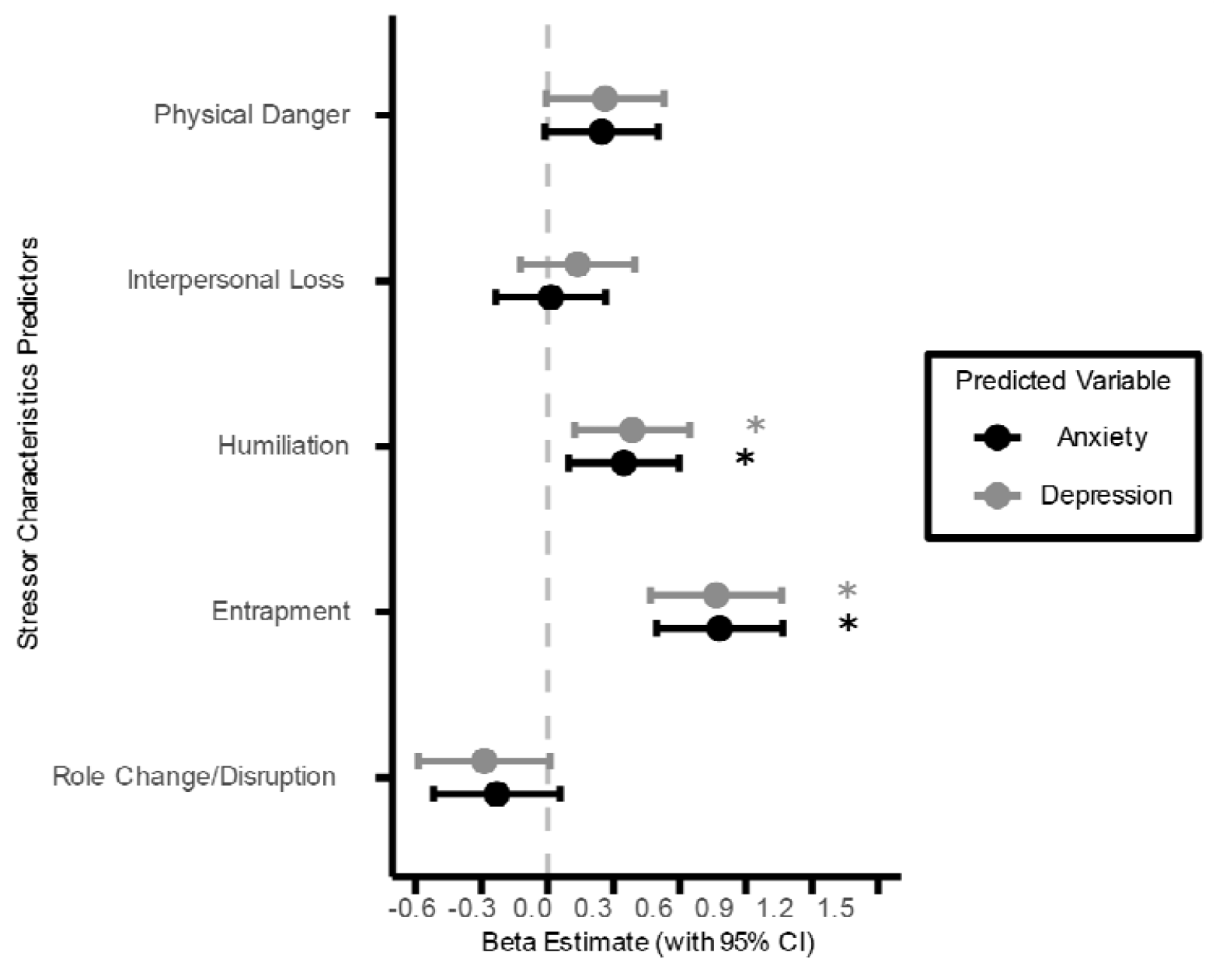
Comparison of standardized β estimates (± 95% confidence intervals) from LME models predicting anxiety and depression symptoms at three time points from total lifetime frequency of five stressor characteristics at baseline assessment. The vertical dashed line represents β = 0. All LME models include a random intercept for each participant and adjust for potentially confounding covariates. Positive β estimates indicate that greater stressor frequency is associated with higher symptom scores. *: q_FDR_ < 0.05

Results from LME models assessing total lifetime severity of the five stressor characteristics as predictors showed similar patterns. After accounting for lifetime severity of four other stressor characteristics and other potential confounding covariates, greater humiliation and entrapment severity at baseline assessment prospectively predicted higher anxiety symptoms (Humiliation: *β* = 0.47, *β*_*back-transformed*_ = 0.25, *p*_*uncorrected*_ = 0.0083, *p*_*FDR*_ = 0.021; Entrapment: *β* = 0.62, *β*_*back-transformed*_ = 0.33, *p*_*uncorrected*_ = 0.0019, *p*_*FDR*_ = 0.0095, **Figure 3**). Although the prospective associations between the lifetime severity of both stressor characteristics and depression symptoms were also positive and significant before correcting for multiple comparisons, only the main effects of entrapment severity survived FDR corrections (Humiliation: *β* = 0.39, *β*_*back-transformed*_ = 0.20, *p*_*uncorrected*_ = 0.035, *p*_*FDR*_ = 0.088; Entrapment: *β* = 0.86, *β*_*back-transformed*_ = 0.44, *p*_*uncorrected*_ = 5.76 × 10^−5^, *p*_*FDR*_ = 0.00029, **Figure 3**).

**Figure 3.**
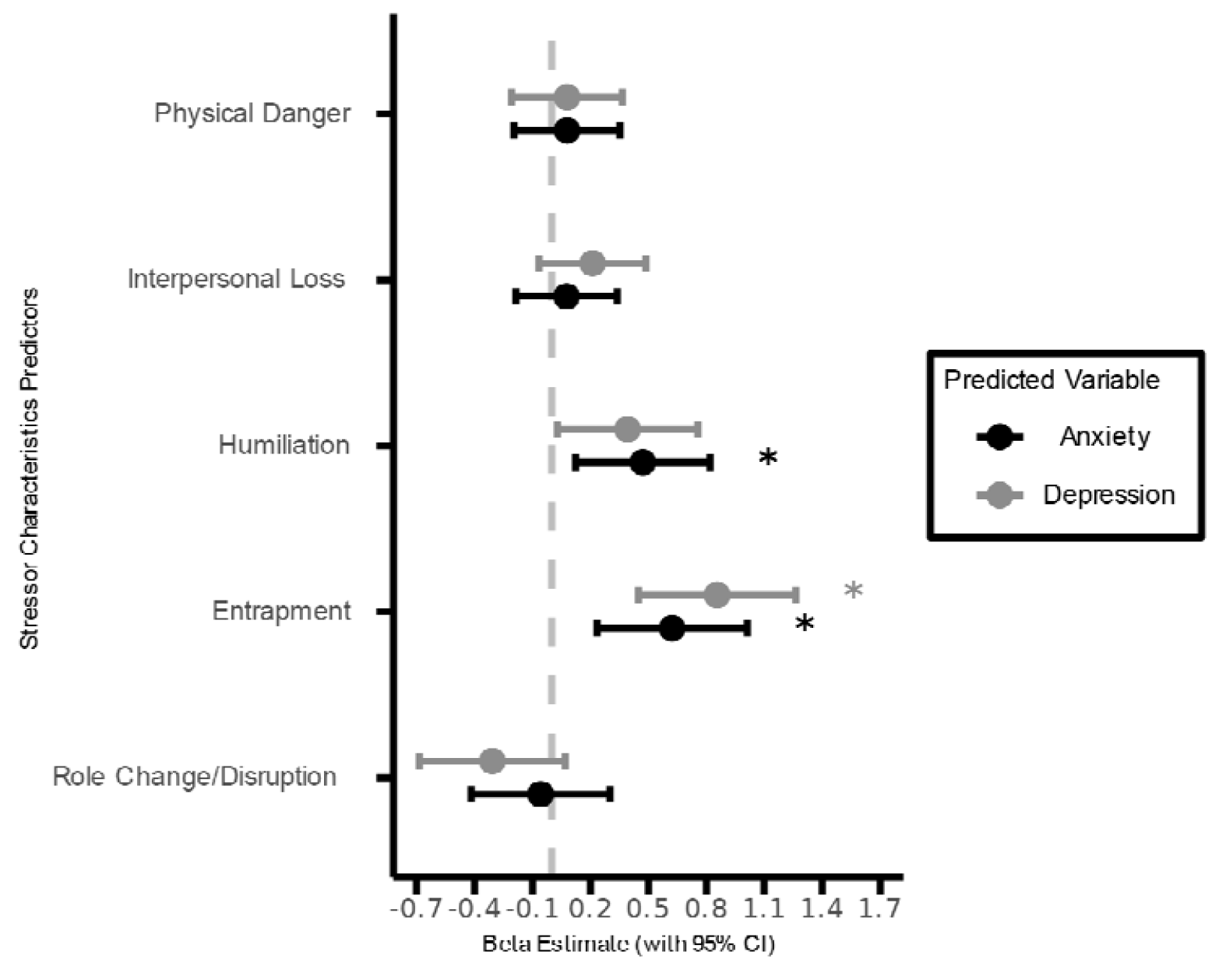
Comparison of standardized β estimates (± 95% confidence intervals) from LME models predicting anxiety and depression symptoms at three time points from total lifetime severity of five stressor characteristics at baseline assessment. The vertical dashed line represents β = 0. All LME models include a random intercept for each participant and adjust for potentially confounding covariates. Positive β estimates indicate that greater stressor severity is associated with higher symptom scores. *: q_FDR_ < 0.05

Overall, these results show that both self-reported frequency and severity of exposures to humiliation and entrapment stressors predict prospective anxiety and depression symptoms at all three time points, suggesting that humiliation and entrapment may be the stressor characteristics most strongly related to anxiety and depression in adolescence.

### Total lifetime frequency and severity of entrapment differentially predict prospective anxiety and depression symptoms

We further tested if total lifetime frequency of any stressor social-psychological characteristic differentially predicted each symptom at three time points by including the other symptom measure at baseline assessment as an additional covariate in each LME model. We found that total lifetime frequency of entrapment stressors differentially predicted anxiety symptoms at all three time points (*β* = 0.56, *β*_*back-transformed*_ = 0.30, *p*_*uncorrected*_ = 7.63 × 10^−5^, *p*_*FDR*_ = 0.00038, **Supplementary Figure 13**), but not depression symptoms (*β* = 0.37, *β*_*back-transformed*_ = 0.19, *p*_*uncorrected*_ = 0.010, *p*_*FDR*_ = 0.052, **Supplementary Figure 13**).

We then conducted similar analyses using total lifetime severity of the five stressor characteristics as differential predictors of prospective anxiety and depression symptoms at three time points. After accounting for baseline depression symptoms, the main effects of humiliation and entrapment severity predicting prospective anxiety symptoms were not significant (Humiliation: *β* = 0.29, *β*_*back-transformed*_ = 0.15, *p*_*uncorrected*_ = 0.086; Entrapment: *β* = 0.34, *β*_*back-transformed*_ = 0.18, *p*_*uncorrected*_ = 0.075, **Supplementary Figure 14**). On the contrary, greater total lifetime severity of entrapment differentially predicted higher depression symptoms after including baseline anxiety symptoms as a covariate (Entrapment: *β* = 0.50, *β*_*back-transformed*_ = 0.26, *p*_*uncorrected*_ = 0.0087, *p*_*FDR*_ = 0.043, **Supplementary Figure 14**).

These results suggest that exposures to entrapment stressors may be a shared risk factor for both anxiety and depression in adolescence, but whether it is differentially associated with one or the other outcome depends on the measure used to assess such exposures. The number of entrapment exposures differentially predicted prospective anxiety symptoms, whereas the subjective severity rating of entrapment exposures differentially predicted prospective depression symptoms

### RSFC between frontoparietal, default and ventral attention networks does not predict prospective anxiety and depression symptoms

Finally, we assessed if RSFC within and between default, frontoparietal and ventral attention networks at baseline predicted each symptom (i.e., anxiety, depression) at each of the three time points (baseline, 6-month and 12-month follow-up assessments). These RSFC metrics were entered as separate predictors (see **Materials and Methods**). After accounting for potential confounding covariates, more positive/less negative RSFC within the default network (*β* = 0.30, *β*_*back-transformed*_ = 0.16, *p*_*uncorrected*_ = 0.040, *p*_*FDR*_ = 0.079, **Figure 4**), between frontoparietal and default networks (*β* = 0.38, *β*_*back-transformed*_ = 0.21, *p*_*uncorrected*_ = 0.0097, *p*_*FDR*_ = 0.058, **Figure 2**), and between frontoparietal and ventral attention networks (*β* = 0.31, *β*_*back-transformed*_ = 0.17, *p*_*uncorrected*_ = 0.034, *p*_*FDR*_ = 0.079, **Figure 4**) predict higher prospective anxiety symptoms. However, the main effects of these associations failed to survive FDR corrections. All prospective associations between RSFC and depression symptoms did not reach statistical significance (*p*_*uncorrected*_ > 0.05, **Figure 4**).

**Figure 4.**
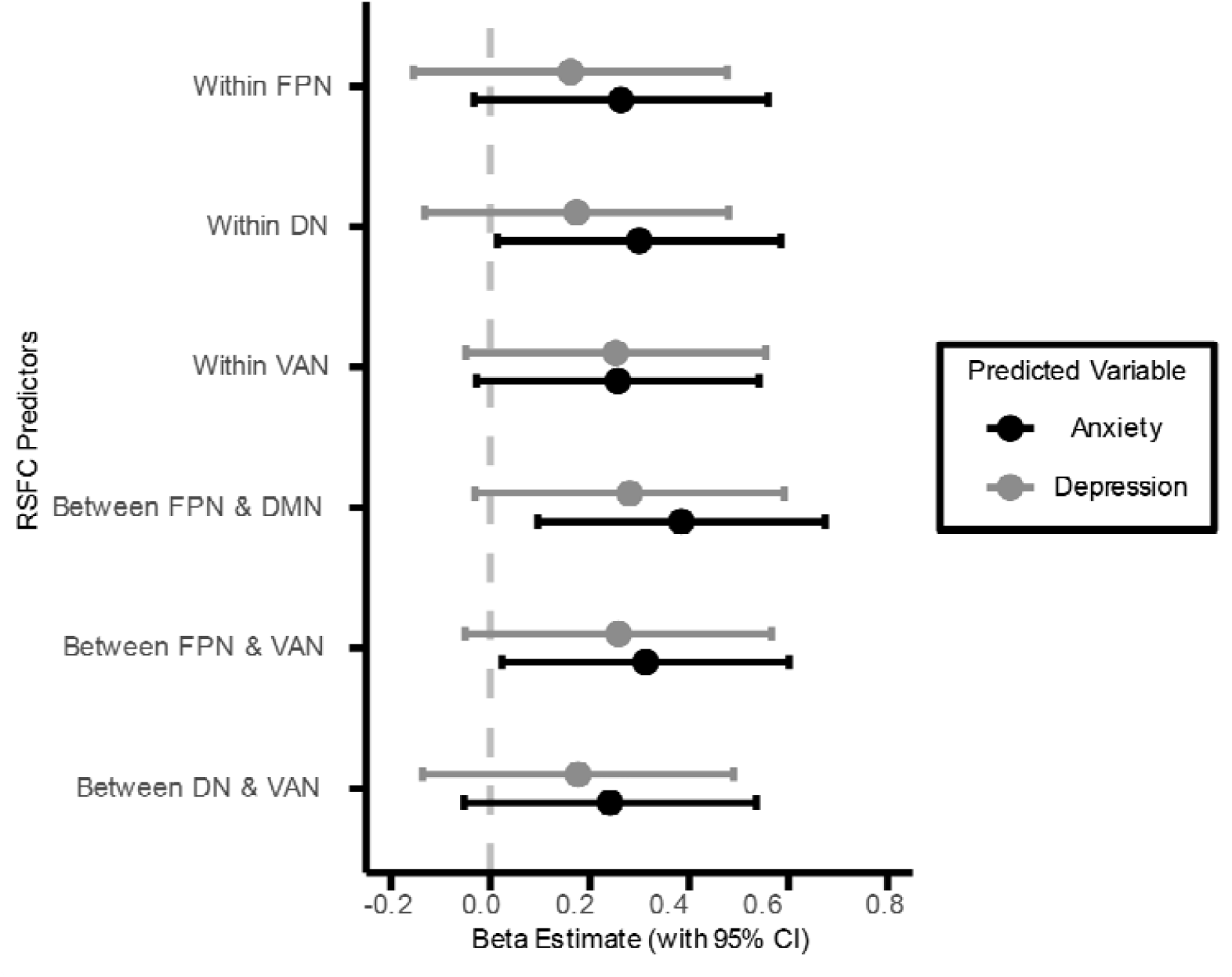
Comparison of standardized β estimates (± 95% confidence intervals) from LME models predicting anxiety and depression symptoms at three time points from RSFC at baseline assessment. Six RSFC metrics are listed on the y-axis: within-network connectivity of the frontoparietal (FPN), default (DN), and ventral attention (VAN) networks, and between-network connectivity of FPN–DN, FPN–VAN, and DN–VAN. The vertical dashed line represents β = 0. All LME models include a random intercept for each participant and adjust for potentially confounding covariates. Positive β estimates indicate that greater RSFC is associated with higher symptom scores. *: q_FDR_ < 0.05

We further assessed if RSFC within and between default, frontoparietal and ventral attention networks at baseline differentially predicted each symptom by accounting for the other symptom measure at baseline assessment in each LME model. As before, RSFC metrics were entered as separate predictors (see **Materials and Methods**). After accounting for potential confounding covariates and depression symptoms at time of baseline assessment, more positive/less negative RSFC between frontoparietal and default networks differentially predict higher prospective anxiety symptoms (*β* = 0.22, *β*_*back-transformed*_ = 0.12, *p*_*uncorrected*_ = 0.046, *p*_*FDR*_ = 0.14, **Supplemental Figure 15**). All other differential associations between RSFC and either symptom did not reach statistical significance (*p*_*uncorrected*_ > 0.05, **Supplemental Figure 15**). Taken together, our results suggest that RSFC between the frontoparietal, default, and ventral attention networks have limited predictive value for future anxiety and depression symptoms and do not differentiate between the two symptom outcomes.

## Discussion

Despite a wealth of prior research examining associations between stress exposures, neurobiology and internalizing psychopathology, it remains unknown if exposures to major life stressors with specific characteristics and exhibiting certain resting-state functional connectivity patterns predict future anxiety and depression symptoms in adolescence. To investigate, we acquired functional neuroimaging at baseline and tracked a sample of adolescents longitudinally for one year, more than half of whom were currently diagnosed with at least one depressive or anxiety disorder at time of initial assessment. We first tested if lifetime frequency and severity of exposures to five stressor characteristics at the time of baseline assessment predicted each symptom (anxiety, depression) at each of the three time points (baseline, 6-month and 12-month follow-up assessments). We then assessed if RSFC within and between the frontoparietal, default, and ventral attention networks at the time of baseline assessment predicted each symptom at the three time points.

Results from the LME models revealed that (a) higher total lifetime frequency and severity of humiliation and entrapment exposures at baseline predicted higher prospective anxiety symptoms; (b) higher total lifetime frequency of humiliation, total lifetime frequency and severity of entrapment exposures at baseline predicted higher prospective depression symptoms; (c) higher total lifetime frequency of entrapment exposure at baseline differentially predicted higher prospective anxiety symptoms above and beyond baseline depression symptoms; (d) higher total lifetime severity of entrapment exposure at baseline differentially predicted higher prospective depression symptoms above and beyond baseline anxiety symptoms; and (e) RSFC within and between frontoparietal, default, and ventral attention networks did not predict either anxiety or depression symptoms. These findings identify humiliation and entrapment as key stressor characteristics predicting future anxiety and depression symptoms in adolescence, while also suggesting that functional interactions among the frontoparietal, default, and ventral attention networks offer limited predictive value for these outcomes.

As opposed to prior studies suggesting that exposure to major life stressors characterized by danger is a specific risk factor for anxiety (Asselmann et al., 2015; Ayazi et al., 2014; Finlay-Jones & Brown, 1981; Kendler et al., 2003), we failed to find a significant association between lifetime exposures to physical danger and prospective anxiety symptoms. Moreover, our results did not reveal differential prediction of depressive symptoms by lifetime exposures to loss stressors, as has been previously shown (Asselmann et al., 2015; Farmer & McGuffin, 2003; Finlay-Jones & Brown, 1981; Keller et al., 2007; Kendler et al., 2003). Unlike prior studies that conducted bivariate analyses between specific stressor characteristics and mental health outcomes, our study examined the *unique contributions* of each stressor characteristics to the prediction of anxiety and depression symptoms after accounting for all other stressor characteristics. Taken together, these results suggest that the differential association between physical danger/interpersonal loss and prospective anxiety/depression symptoms may be fully explained by the main effects of exposures to other stressor characteristics.

Our results are consistent with prior studies implying that exposures to humiliation and entrapment may contribute to both anxiety and depression (Griffiths et al., 2014; Kendler et al., 2003; Li et al., 2024; Taylor et al., 2011). Humiliation is theorized as perceived devaluation of the self arising from being unfairly harmed by others (Klein, 1991; D. A. Lee et al., 2001).

Exposures to humiliation may heighten anxiety and depression by increasing vigilance towards potential future harm by others and viewing the self as worthless. Common examples of humiliation stressors include public shaming, peer bullying, social exclusion, rejection and revelation of infidelity. We found that humiliation frequency predicted both anxiety and depression symptoms over time, but differentially predicted neither. Humiliation severity at baseline predicted prospective anxiety symptoms, while the prospective associations between humiliation severity and depression symptoms failed to survive FDR corrections. On the other hand, entrapment refers to the feelings of being chronically trapped in a stressful situation despite having a strong desire to escape (Gilbert & Allan, 1998). Exposures to entrapment stressors may provoke anxiety and depression by fostering a sense of diminished control and reinforcing perceptions of futility, consistent with the concept of learned helplessness. Examples of entrapment stressors include persistent food and financial insecurity, unsafe housing and overwhelming demands at work or school. We found that entrapment frequency and severity predicted both anxiety and depression symptoms over time. Results from the present study highlight the potentially long-lasting impacts of humiliating and socio-structural stressors on both anxiety and depression. Another interesting observation from our findings is that frequency and severity measures of entrapment exposures are differentially associated with anxiety and depression symptoms respectively. This suggests that more frequent entrapment exposures, irrespective of severity, may be more anxio-genic by encouraging hypervigilance sustaining anticipation of further threat. On the other hand, isolated occurrences of severe entrapment stressors can exhaust one’s coping resources and perpetrate the belief that any escape is futile.

Finally, we uncovered nominally significant associations between RSFC within the default network, between frontoparietal and default networks, and between frontoparietal and ventral attention networks and prospective anxiety symptoms. Our results are not consistent with the formulation that abnormal interactions between functional networks theorized to support executive control (i.e. frontoparietal), self-referential processing (i.e. default) and salience processing (i.e. ventral attention) are associated with anxiety and depression (Al-Ezzi et al., 2021; Berman et al., 2011; Duan et al., 2020; Ho et al., 2015; Kaiser et al., 2015; Leech & Sharp, 2014; Manoliu et al., 2014; Nejad et al., 2013; Pawlak et al., 2022; Perino et al., 2021; Rzepa & McCabe, 2018; Sievertsen et al., 2025; Willinger et al., 2024; Y. Zhang et al., 2024). The lack of significant associations may arise from methodological differences compared to the prior literature. For example, while our study examined prospective associations between RSFC and affective symptoms, most previous studies focused on cross-sectional associations (Al-Ezzi et al., 2021; Ho et al., 2015; Kaiser et al., 2015; Manoliu et al., 2014; Perino et al., 2021; Rzepa & McCabe, 2018, 2018; Willinger et al., 2024; Y. Zhang et al., 2024). Among the longitudinal studies, most focused on functional connectivity involving isolated network ROIs, especially subcortical ROIs, as predictors of anxiety and depression symptoms at follow-up (Connolly et al., 2017; Davey et al., 2015; Edalati et al., 2023; Lopez et al., 2018; Pawlak et al., 2022), while we computed RSFC across multiple cortical ROIs for each network of interest (Schaefer et al., 2018). Aside from methodological differences, the lack of significant findings in our study may also be attributed to the limited size of our current sample that may not afford sufficient power for detecting certain prospective brain-behavior associations with small effect sizes (Marek et al., 2022). Lastly, our study did not find differential associations of any RSFC within and between the frontoparietal, default and ventral attention networks with respect to either anxiety or depression symptoms. These results suggest that there may not be any symptom-specific RSFC predictor.

### Strengths and Limitations

Several strengths and limitations of this work should be noted. In terms of strengths, we leveraged a longitudinal sample of adolescents enriched for clinical diagnosis of anxiety and depressive disorders with a narrow age range. This enabled us to examine prospective associations between exposure to major life stressors, neurobiology, and clinical symptoms during the developmental stage characterized by heightened vulnerability to stress-related psychopathology (Casey et al., 2008; Larsen & Luna, 2018; March-Llanes et al., 2017; Merikangas et al., 2010; Paus et al., 2008). We assessed a rich repertoire of theoretically relevant stressors occurring across the entire life course using distinct dimensions of social-psychological characteristics assessed by the STRAIN. Our study had a novel focus on using subjective severity ratings besides the number of exposures to stressor characteristics. In this way, we demonstrated the validity of these stressor characteristic indices in predicting internalizing psychopathology over time. We included all five social-psychological characteristics and the potential confounding variables into the same LME model when predicting anxiety/depression symptoms at three time points. This ensured that any prospective association between a stressor characteristic and clinical symptom is robust to the inclusion of other inter-correlated stressor characteristics. Lastly, each participant in our study underwent 23.2 minutes of rsfMRI. The scan duration of our rsfMRI data goes far beyond that of most previous rsfMRI studies and should afford relatively high reliability (Birn et al., 2013).

Several limitations should also be noted. First, the sample was relatively small (*n* = 150), which did not afford us sufficient statistical power to assess RSFC involving other functional networks such as the dorsal attention and the limbic networks as well as the subcortical structures such as the amygdala, which have also been associated with anxiety and depression (Brandl et al., 2022; Connolly et al., 2017; Cullen et al., 2014; Kaiser et al., 2015; Liu et al., 2021; Sylvester et al., 2012; Xu et al., 2019; Z. Zhang et al., 2025). Future high-powered analyses would benefit from a whole-brain approach. Although small sample size is a common issue with richly-phenotyped, longitudinal datasets involving clinical samples, findings from the present study should be validated by larger samples in future studies. Since the fMRI protocol of the current dataset was harmonized with other HCP studies (Siless et al., 2020), other HCP datasets may provide valuable resources for this purpose. Second, the vast majority (79.33%) of participants were White, came from highly educated families, and endorsed relatively low exposures to stressors characterized by danger and role change/disruption (**Table 1**) compared to previous studies on autistic adults (Moseley et al., 2025) and sexual minority adults of color (Parra et al., 2023). Although our study assessed moderate-to-severe life stressors, and previous research suggests that exposure to a single severe stressor can trigger depressive episodes in over half of individuals (Monroe et al., 2007; Shapiro, 1979), further research is still needed to confirm whether our findings generalize to populations with greater exposure to stressors involving danger and significant role disruptions. Lastly, although we included a number of covariates such as age, race and ethnicity, baseline symptoms, and diagnostic status in our LME models to ensure that our findings could not be explained by these confounders, other potential confounders such as medication use and family history of psychopathology were unavailable in this dataset and may be relevant.

## Conclusion

Notwithstanding these limitations, through the use a longitudinal sample of adolescents, over half of whom met clinical cut-offs for at least one anxiety or depressive disorder, the present study demonstrate that lifetime frequency and severity of humiliation and entrapment exposures predict both anxiety and depression symptoms at baseline, 6 months, and 12 months later. These results imply that among all characteristics of major life stressors examined by the Adolescent STRAIN, humiliation and entrapment may be the most important predictive characteristics of both anxiety and depression over time. Follow-up analyses showed that lifetime frequency of entrapment exposures differentially predicts anxiety symptoms, whereas lifetime severity differentially predicts depression symptoms, suggesting that distinct measures of stressor exposures may be associated with different mental health outcomes. Finally, our results showed no association between RSFC among the frontoparietal, default, and ventral attention networks and anxiety or depression symptoms across all three time points, indicating that these connectivity patterns may be relatively weak predictors of adolescent internalizing symptoms. Looking forward, additional research is needed to elucidate how lifetime stressor exposure interacts with brain activity and connectivity—as well as peripheral biological processes—to predict other forms of psychopathology in adolescence and across the life course.

## References

Al-Ezzi, A., Kamel, N., Faye, I., & Gunaseli, E. (2021). Analysis of Default Mode Network in Social Anxiety Disorder: EEG Resting-State Effective Connectivity Study. Sensors, 21(12), Article 12. 10.3390/s21124098

American Psychiatric Association. (2013). Diagnostic and Statistical Manual of Mental Disorders (5th ed.). American Psychiatric Association. https://psychiatryonline.org/doi/epub/10.1176/appi.books.9780890425596

Angold, A., Costello, E. J., Messer, S. C., & Pickles, A. (1995). Development of a short questionnaire for use in epidemiological studies of depression in children and adolescents. International Journal of Methods in Psychiatric Research, 5(4), 237–249.

Asselmann, E., Wittchen, H.-U., Lieb, R., Höfler, M., & Beesdo-Baum, K. (2015). Danger and loss events and the incidence of anxiety and depressive disorders: A prospective-longitudinal community study of adolescents and young adults. Psychological Medicine, 45(1), 153–163. 10.1017/S0033291714001160

Ayazi, T., Lien, L., Eide, A., Swartz, L., & Hauff, E. (2014). Association between exposure to traumatic events and anxiety disorders in a post-conflict setting: A cross-sectional community study in South Sudan. BMC Psychiatry, 14, 6. 10.1186/1471-244X-14-6

Bates, D., Mächler, M., Bolker, B. M., & Walker, S. C. (2015). Fitting linear mixed-effects models using lme4. Journal of Statistical Software, 67(1), 1–48. 10.18637/jss.v067.i01

Berman, M. G., Peltier, S., Nee, D. E., Kross, E., Deldin, P. J., & Jonides, J. (2011). Depression, rumination and the default network. Social Cognitive and Affective Neuroscience, 6(5), 548–555. 10.1093/scan/nsq080

Birn, R. M., Molloy, E. K., Patriat, R., Parker, T., Meier, T. B., Kirk, G. R., Nair, V. A., Meyerand, M. E., & Prabhakaran, V. (2013). The effect of scan length on the reliability of resting-state fMRI connectivity estimates. NeuroImage, 83, 550–558. 10.1016/j.neuroimage.2013.05.099

Bonanno, G. A., Chen, S., & Galatzer-Levy, I. R. (2023). Resilience to potential trauma and adversity through regulatory flexibility. Nature Reviews Psychology, 2(11), 663–675. 10.1038/s44159-023-00233-5

Brandl, F., Weise, B., Mulej Bratec, S., Jassim, N., Hoffmann Ayala, D., Bertram, T., Ploner, M., & Sorg, C. (2022). Common and specific large-scale brain changes in major depressive disorder, anxiety disorders, and chronic pain: A transdiagnostic multimodal meta-analysis of structural and functional MRI studies. Neuropsychopharmacology, 47(5), 1071–1080. 10.1038/s41386-022-01271-y

Brown, G. W., Harris, T. O., & Hepworth, C. (1995). Loss, humiliation and entrapment among women developing depression: A patient and non-patient comparison. Psychological Medicine, 25(1), 7–21. 10.1017/s003329170002804x

Burleson Daviss, W., Birmaher, B., Melhem, N. A., Axelson, D. A., Michaels, S. M., & Brent, D. A. (2006). Criterion validity of the Mood and Feelings Questionnaire for depressive episodes in clinic and non-clinic subjects. Journal of Child Psychology and Psychiatry, 47(9), 927–934. 10.1111/j.1469-7610.2006.01646.x

Casey, B. j., Jones, R. M., & Hare, T. A. (2008). The Adolescent Brain. Annals of the New York Academy of Sciences, 1124(1), 111–126. 10.1196/annals.1440.010

Chorpita, B. F., Yim, L., Moffitt, C., Umemoto, L. A., & Francis, S. E. (2000). Assessment of symptoms of DSM-IV anxiety and depression in children: A revised child anxiety and depression scale. Behaviour Research and Therapy, 38(8), 835–855. 10.1016/S0005-7967(99)00130-8

Cohen, S., Murphy, M. L. M., & Prather, A. A. (2019). Ten Surprising Facts About Stressful Life Events and Disease Risk. Annual Review of Psychology, 70, 577–597. 10.1146/annurev-psych-010418-102857

Cole, M. W., Repovš, G., & Anticevic, A. (2014). The frontoparietal control system: A central role in mental health. Neuroscientist, 20(6), 652–664. 10.1177/1073858414525995

Connolly, C. G., Ho, T. C., Blom, E. H., LeWinn, K. Z., Sacchet, M. D., Tymofiyeva, O., Simmons, A. N., & Yang, T. T. (2017). Resting-state functional connectivity of the amygdala and longitudinal changes in depression severity in adolescent depression. Journal of Affective Disorders, 207, 86–94. 10.1016/j.jad.2016.09.026

Costello, E. J., & Angold, A. (1988). Scales to Assess Child and Adolescent Depression: Checklists, Screens, and Nets. Journal of the American Academy of Child & Adolescent Psychiatry, 27(6), 726–737. 10.1097/00004583-198811000-00011

Cullen, K. R., Westlund, M. K., Klimes-Dougan, B., Mueller, B. A., Houri, A., Eberly, L. E., & Lim, K. O. (2014). Abnormal Amygdala Resting-State Functional Connectivity in Adolescent Depression. JAMA Psychiatry, 71(10), 1138–1147. 10.1001/jamapsychiatry.2014.1087

Davey, C. G., Whittle, S., Harrison, B. J., Simmons, J. G., Byrne, M. L., Schwartz, O. S., & Allen, N. B. (2015). Functional brain-imaging correlates of negative affectivity and the onset of first-episode depression. Psychological Medicine, 45(5), 1001–1009. 10.1017/S0033291714002001

Duan, L., Van Dam, N. T., Ai, H., & Xu, P. (2020). Intrinsic organization of cortical networks predicts state anxiety: An functional near-infrared spectroscopy (fNIRS) study. Translational Psychiatry, 10(1), 402. 10.1038/s41398-020-01088-7

Edalati, H., Afzali, M. H., Spinney, S., Bourque, J., Dagher, A., & Conrod, P. J. (2023). A longitudinal mediation study of peer victimization and resting-state functional connectivity as predictors of development of adolescent psychopathology. Frontiers in Psychiatry, 14. 10.3389/fpsyt.2023.1099772

Faravelli, C., & Pallanti, S. (1989). Recent life events and panic disorder. The American Journal of Psychiatry, 146(5), 622–626. 10.1176/ajp.146.5.622

Farmer, A. E., & McGuffin, P. (2003). Humiliation, loss and other types of life events and difficulties: A comparison of depressed subjects, healthy controls and their siblings. Psychological Medicine, 33(7), 1169–1175. 10.1017/s0033291703008419

Finlay-Jones, R., & Brown, G. W. (1981). Types of stressful life event and the onset of anxiety and depressive disorders. Psychological Medicine, 11(4), 803–815. 10.1017/S0033291700041301

Francis, J. L., Moitra, E., Dyck, I., & Keller, M. B. (2012). The impact of stressful life events on relapse of generalized anxiety disorder. Depression and Anxiety, 29(5), 386–391. 10.1002/da.20919

Gilbert, P., & Allan, S. (1998). The role of defeat and entrapment (arrested flight) in depression: An exploration of an evolutionary view. Psychological Medicine, 28(3), 585–598. 10.1017/S0033291798006710

Glasser, M. F., Smith, S. M., Marcus, D. S., Andersson, J. L. R., Auerbach, E. J., Behrens, T. E. J., Coalson, T. S., Harms, M. P., Jenkinson, M., Moeller, S., Robinson, E. C., Sotiropoulos, S. N., Xu, J., Yacoub, E., Ugurbil, K., & Van Essen, D. C. (2016). The Human Connectome Project’s neuroimaging approach. Nature Neuroscience, 19(9), 1175–1187. 10.1038/nn.4361

Glasser, M. F., Sotiropoulos, S. N., Wilson, J. A., Coalson, T. S., Fischl, B., Andersson, J. L., Xu, J., Jbabdi, S., Webster, M., Polimeni, J. R., Van Essen, D. C., & Jenkinson, M. (2013). The minimal preprocessing pipelines for the Human Connectome Project. NeuroImage, 80, 105–124. 10.1016/J.NEUROIMAGE.2013.04.127

Greve, D. N., & Fischl, B. (2009). Accurate and Robust Brain Image Alignment using Boundary-based Registration. NeuroImage, 48(1), 63–72. 10.1016/j.neuroimage.2009.06.060

Griffanti, L., Salimi-Khorshidi, G., Beckmann, C. F., Auerbach, E. J., Douaud, G., Sexton, C. E., Zsoldos, E., Ebmeier, K. P., Filippini, N., Mackay, C. E., Moeller, S., Xu, J., Yacoub, E., Baselli, G., Ugurbil, K., Miller, K. L., & Smith, S. M. (2014). ICA-based artefact removal and accelerated fMRI acquisition for improved resting state network imaging. NeuroImage, 95, 232–247. 10.1016/j.neuroimage.2014.03.034

Griffiths, A. W., Wood, A. M., Maltby, J., Taylor, P. J., & Tai, S. (2014). The prospective role of defeat and entrapment in depression and anxiety: A 12-month longitudinal study. Psychiatry Research, 216(1), 52–59. 10.1016/j.psychres.2014.01.037

Ho, T. C., Connolly, C. G., Henje Blom, E., LeWinn, K. Z., Strigo, I. A., Paulus, M. P., Frank, G., Max, J. E., Wu, J., Chan, M., Tapert, S. F., Simmons, A. N., & Yang, T. T. (2015). Emotion-Dependent Functional Connectivity of the Default Mode Network in Adolescent Depression. Biological Psychiatry, 78(9), 635–646. 10.1016/j.biopsych.2014.09.002

Huang, P., Chan, S. Y., Ngoh, Z. M., Nadarajan, R., Chong, Y. S., Gluckman, P. D., Chen, H., Fortier, M. V., Tan, A. P., & Meaney, M. J. (2023). Functional connectivity analysis of childhood depressive symptoms. NeuroImage: Clinical, 38, 103395. 10.1016/j.nicl.2023.103395

Hubbard, N. A., Bauer, C. C. C., Siless, V., Auerbach, R. P., Elam, J. S., Frosch, I. R., Henin, A., Hofmann, S. G., Hodge, M. R., Jones, R., Lenzini, P., Lo, N., Park, A. T., Pizzagalli, D. A., Vaz-DeSouza, F., Gabrieli, J. D. E., Whitfield-Gabrieli, S., Yendiki, A., & Ghosh, S. S. (2024). The Human Connectome Project of adolescent anxiety and depression dataset. Scientific Data, 11(1), 837. 10.1038/s41597-024-03629-x

Jalbrzikowski, M., Larsen, B., Hallquist, M. N., Foran, W., Calabro, F., & Luna, B. (2017). Development of White Matter Microstructure and Intrinsic Functional Connectivity Between the Amygdala and Ventromedial Prefrontal Cortex: Associations With Anxiety and Depression. Biological Psychiatry, 82(7), 511–521. 10.1016/j.biopsych.2017.01.008

Jenkinson, M., Bannister, P., Brady, M., & Smith, S. (2002). Improved optimization for the robust and accurate linear registration and motion correction of brain images. NeuroImage, 17(2), 825–841. 10.1016/s1053-8119(02)91132-8

Jenkinson, M., & Smith, S. (2001). A global optimisation method for robust affine registration of brain images. Medical Image Analysis, 5(2), 143–156. 10.1016/s1361-8415(01)00036-6

Jin, J., Van Snellenberg, J. X., Perlman, G., DeLorenzo, C., Klein, D. N., Kotov, R., & Mohanty, A. (2020). Intrinsic neural circuitry of depression in adolescent females. Journal of Child Psychology and Psychiatry, 61(4), 480–491. 10.1111/jcpp.13123

Kaiser, R. H., Andrews-Hanna, J. R., Wager, T. D., & Pizzagalli, D. A. (2015). Large-scale network dysfunction in major depressive disorder: A meta-analysis of resting-state functional connectivity. JAMA Psychiatry, 72(6), 603–611. 10.1001/jamapsychiatry.2015.0071

Keller, M. C., Neale, M. C., & Kendler, K. S. (2007). Association of different adverse life events with distinct patterns of depressive symptoms. The American Journal of Psychiatry, 164(10), 1521–1529. 10.1176/APPI.AJP.2007.06091564

Kendler, K. S., Hettema, J. M., Butera, F., Gardner, C. O., & Prescott, C. A. (2003). Life event dimensions of loss, humiliation, entrapment, and danger in the prediction of onsets of major depression and generalized anxiety. Archives of General Psychiatry, 60(8), 789–796. 10.1001/ARCHPSYC.60.8.789

Kendler, K. S., Karkowski, L. M., & Prescott, C. A. (1999). Causal relationship between stressful life events and the onset of major depression. American Journal of Psychiatry, 156(6), 837–841. 10.1176/AJP.156.6.837/ASSET/IMAGES/LARGE/AN5T1.JPEG

Kessler, R. C. (1997). The effects of stressful life events on depression. Annual Review of Psychology, 48, 191–214. 10.1146/annurev.psych.48.1.191

Klein, D. C. (1991). The humiliation dynamic: An overview. Journal of Primary Prevention, 12(2), 93–121. 10.1007/BF02015214

Larsen, B., & Luna, B. (2018). Adolescence as a neurobiological critical period for the development of higher-order cognition. Neuroscience & Biobehavioral Reviews, 94, 179–195. 10.1016/j.neubiorev.2018.09.005

Lee, A. H., Kitagawa, Y., Mirhashem, R., Rodriguez, M., Hilerio, R., & Bernard, K. (2025). Do dimensions of childhood adversity differ in their direct associations with youth psychopathology? A meta-analysis. Development and Psychopathology, 37(2), 871–901. 10.1017/S0954579424000737

Lee, D. A., Scragg, P., & Turner, S. (2001). The role of shame and guilt in traumatic events: A clinical model of shame-based and guilt-based PTSD. British Journal of Medical Psychology, 74(4), 451–466. 10.1348/000711201161109

Leech, R., & Sharp, D. J. (2014). The role of the posterior cingulate cortex in cognition and disease. BrainlJ: A Journal of Neurology, 137(Pt 1), 12–32. 10.1093/BRAIN/AWT162

Li, W. W., Heward, C., Merrick, A., Astridge, B., & Leow, T. (2024). Prevalence of experiencing public humiliation and its effects on victims’ mental health: A systematic review and meta-analysis. Journal of Pacific Rim Psychology, 18, 18344909241252325. 10.1177/18344909241252325

Liu, R., Wang, Y., Chen, X., Zhang, Z., Xiao, L., & Zhou, Y. (2021). Anhedonia correlates with functional connectivity of the nucleus accumbens subregions in patients with major depressive disorder. NeuroImagelJ: Clinical, 30, 102599. 10.1016/J.NICL.2021.102599

Lopez, K. C., Luby, J. L., Belden, A. C., & Barch, D. M. (2018). Emotion dysregulation and functional connectivity in children with and without a history of major depressive disorder. Cognitive, Affective, & Behavioral Neuroscience, 18(2), 232–248. 10.3758/s13415-018-0564-x

Manoliu, A., Meng, C., Brandl, F., Doll, A., Tahmasian, M., Scherr, M., Schwerthöffer, D., Zimmer, C., Förstl, H., Bäuml, J., Riedl, V., Wohlschläger, A. M., & Sorg, C. (2014). Insular dysfunction within the salience network is associated with severity of symptoms and aberrant inter-network connectivity in major depressive disorder. Frontiers in Human Neuroscience, 7(JAN). 10.3389/fnhum.2013.00930

March-Llanes, J., Marqués-Feixa, L., Mezquita, L., Fañanás, L., & Moya-Higueras, J. (2017). Stressful life events during adolescence and risk for externalizing and internalizing psychopathology: A meta-analysis. European Child & Adolescent Psychiatry, 26(12), 1409–1422. 10.1007/s00787-017-0996-9

Marek, S., Tervo-Clemmens, B., Calabro, F. J., Montez, D. F., Kay, B. P., Hatoum, A. S., Donohue, M. R., Foran, W., Miller, R. L., Hendrickson, T. J., Malone, S. M., Kandala, S., Feczko, E., Miranda-Dominguez, O., Graham, A. M., Earl, E. A., Perrone, A. J., Cordova, M., Doyle, O., … Dosenbach, N. U. F. (2022). Reproducible brain-wide association studies require thousands of individuals. Nature 2022 603:7902, 603(7902), 654–660. 10.1038/s41586-022-04492-9

Menon, V. (2011). Large-scale brain networks and psychopathology: A unifying triple network model. Trends in Cognitive Sciences, 15(10), 483–506. 10.1016/j.tics.2011.08.003

Merikangas, K. R., He, J., Burstein, M., Swanson, S. A., Avenevoli, S., Cui, L., Benjet, C., Georgiades, K., & Swendsen, J. (2010). Lifetime Prevalence of Mental Disorders in U.S. Adolescents: Results from the National Comorbidity Survey Replication–Adolescent Supplement (NCS-A). Journal of the American Academy of Child & Adolescent Psychiatry, 49(10), 980–989. 10.1016/j.jaac.2010.05.017

Miloyan, B., Bienvenu, O. J., Brilot, B., & Eaton, W. W. (2018). Adverse life events and the onset of anxiety disorders. Psychiatry Research, 259, 488–492. 10.1016/j.psychres.2017.11.027

Monroe, S. M., Slavich, G. M., Torres, L. D., & Gotlib, I. H. (2007). Major life events and major chronic difficulties are differentially associated with history of major depressive episodes. Journal of Abnormal Psychology, 116(1), 116–124. 10.1037/0021-843X.116.1.116

Moseley, R. L., Hedley, D., Gamble-Turner, J. M., Uljarević, M., Bury, S. M., Shields, G. S., Trollor, J. N., Stokes, M. A., & Slavich, G. M. (2025). Lifetime stressor exposure is related to suicidality in autistic adults: A multinational study. Autism, 29(5), 1184–1208. 10.1177/13623613241299872

Nejad, A. B., Fossati, P., & Lemogne, C. (2013). Self-referential processing, rumination, and cortical midline structures in major depression. Frontiers in Human Neuroscience, 7(OCT). 10.3389/fnhum.2013.00666

Parkes, L., Fulcher, B., Yücel, M., & Fornito, A. (2018). An evaluation of the efficacy, reliability, and sensitivity of motion correction strategies for resting-state functional MRI. NeuroImage, 171, 415–436. 10.1016/j.neuroimage.2017.12.073

Parra, L. A., Spahr, C. M., Goldbach, J. T., Bray, B. C., Kipke, M. D., & Slavich, G. M. (2023). Greater lifetime stressor exposure is associated with poorer mental health among sexual minority people of color. Journal of Clinical Psychology, 79(4), 1130–1155. 10.1002/jclp.23463

Paus, T., Keshavan, M., & Giedd, J. N. (2008). Why do many psychiatric disorders emerge during adolescence? Nature Reviews Neuroscience, 9(12), 947–957. 10.1038/nrn2513

Pawlak, M., Bray, S., & Kopala-Sibley, D. C. (2022). Resting state functional connectivity as a marker of internalizing disorder onset in high-risk youth. Scientific Reports, 12(1), 21337. 10.1038/s41598-022-25805-y

Perino, M. T., Myers, M. J., Wheelock, M. D., Yu, Q., Harper, J. C., Manhart, M. F., Gordon, E. M., Eggebrecht, A. T., Pine, D. S., Barch, D. M., Luby, J. L., & Sylvester, C. M. (2021). Whole-Brain Resting-State Functional Connectivity Patterns Associated With Pediatric Anxiety and Involuntary Attention Capture. Biological Psychiatry Global Open Science, 1(3), 229–238. 10.1016/j.bpsgos.2021.05.007

Power, J. D., Barnes, K. A., Snyder, A. Z., Schlaggar, B. L., & Petersen, S. E. (2012). Spurious but systematic correlations in functional connectivity MRI networks arise from subject motion. NeuroImage, 59(3), 2142–2154. 10.1016/J.NEUROIMAGE.2011.10.018

Qu, Y., Rappaport, B. I., Luby, J. L., & Barch, D. M. (2021). No associations in preregistered study of youth depression and functional connectivity of fronto-parietal and default mode networks. Neuroimage: Reports, 1(3), 100036. 10.1016/j.ynirp.2021.100036

Robinson, E. C., Jbabdi, S., Glasser, M. F., Andersson, J., Burgess, G. C., Harms, M. P., Smith, S. M., Van Essen, D. C., & Jenkinson, M. (2014). MSM: A new flexible framework for Multimodal Surface Matching. NeuroImage, 100, 414–426. 10.1016/j.neuroimage.2014.05.069

Rzepa, E., & McCabe, C. (2018). Anhedonia and depression severity dissociated by dmPFC resting-state functional connectivity in adolescents. Journal of Psychopharmacology, 32(10), 1067–1074. 10.1177/0269881118799935

Salimi-Khorshidi, G., Douaud, G., Beckmann, C. F., Glasser, M. F., Griffanti, L., & Smith, S. M. (2014). Automatic denoising of functional MRI data: Combining independent component analysis and hierarchical fusion of classifiers. NeuroImage, 90, 449–468. 10.1016/j.neuroimage.2013.11.046

Satterthwaite, T. D., Elliott, M. A., Gerraty, R. T., Ruparel, K., Loughead, J., Calkins, M. E., Eickhoff, S. B., Hakonarson, H., Gur, R. C., Gur, R. E., & Wolf, D. H. (2013). An improved framework for confound regression and filtering for control of motion artifact in the preprocessing of resting-state functional connectivity data. NeuroImage, 64(1), 240–256. 10.1016/J.NEUROIMAGE.2012.08.052

Schaefer, A., Kong, R., Gordon, E. M., Laumann, T. O., Zuo, X.-N., Holmes, A. J., Eickhoff, S. B., & Yeo, B. T. T. (2018). Local-Global Parcellation of the Human Cerebral Cortex from Intrinsic Functional Connectivity MRI. Cerebral Cortex (New York, N.Y.□: 1991), 28(9), 3095–3114. 10.1093/CERCOR/BHX179

Shapiro, M. B. (1979). The social origins of depression: By G.W. Brown and T. Harris: Its methodological philosophy. Behaviour Research and Therapy, 17(6), 597–603. 10.1016/0005-7967(79)90104-9

Shields, G. S., Fassett-Carman, A., Gray, Z. J., Gonzales, J. E., Snyder, H. R., & Slavich, G. M. (2023). Why is subjective stress severity a stronger predictor of health than stressor exposure? A preregistered two-study test of two hypotheses. Stress and Health, 39(1), 87–102. 10.1002/smi.3165

Sievertsen, S. A., Zhu, J., Fang, A., & Forsyth, J. K. (2025). Resting-State Cortical Network and Subcortical Hyperconnectivity in Youth With Generalized Anxiety Disorder in the ABCD Study. Biological Psychiatry: Cognitive Neuroscience and Neuroimaging. 10.1016/j.bpsc.2025.02.005

Siless, V., Hubbard, N. A., Jones, R., Wang, J., Lo, N., Bauer, C. C. C., Goncalves, M., Frosch, I., Norton, D., Vergara, G., Conroy, K., De Souza, F. V., Rosso, I. M., Wickham, A. H., Cosby, E. A., Pinaire, M., Hirshfeld-Becker, D., Pizzagalli, D. A., Henin, A., … Yendiki, A. (2020). Image acquisition and quality assurance in the Boston Adolescent Neuroimaging of Depression and Anxiety study. NeuroImage□: Clinical, 26, 102242. 10.1016/J.NICL.2020.102242

Slavich, G. M. (2019). Stressnology: The primitive (and problematic) study of life stress exposure and pressing need for better measurement. Brain, Behavior, and Immunity, 75, 3–5. 10.1016/j.bbi.2018.08.011

Slavich, G. M., & Shields, G. S. (2018). Assessing Lifetime Stress Exposure Using the Stress and Adversity Inventory for Adults (Adult STRAIN): An Overview and Initial Validation. Psychosomatic Medicine, 80(1), 17–27. 10.1097/PSY.0000000000000534

Slavich, G. M., Stewart, J. G., Esposito, E. C., Shields, G. S., & Auerbach, R. P. (2019). The Stress and Adversity Inventory for Adolescents (Adolescent STRAIN): Associations with mental and physical health, risky behaviors, and psychiatric diagnoses in youth seeking treatment. Journal of Child Psychology and Psychiatry, 60(9), 998–1009. 10.1111/jcpp.13038

Slotter, E. B., & and Walsh, C. M. (2017). All role transitions are not experienced equally: Associations among self-change, emotional reactions, and self-concept clarity. Self and Identity, 16(5), 531–556. 10.1080/15298868.2017.1280528

Stekhoven, D. J., & Bühlmann, P. (2012). MissForest—Non-parametric missing value imputation for mixed-type data. Bioinformatics, 28(1), 112–118. 10.1093/bioinformatics/btr597

Strikwerda-Brown, C., Davey, C. G., Whittle, S., Allen, N. B., Byrne, M. L., Schwartz, O. S., Simmons, J. G., Dwyer, D., & Harrison, B. J. (2014). Mapping the relationship between subgenual cingulate cortex functional connectivity and depressive symptoms across adolescence. Social Cognitive and Affective Neuroscience, 10(7), 961–968. 10.1093/scan/nsu143

Sund, A. M., Larsson, B., & Wichstrøm, L. (2001). Depressive symptoms among young Norwegian adolescents as measured by The Mood and Feelings Questionnaire (MFQ). European Child & Adolescent Psychiatry, 10(4), 222–229. 10.1007/s007870170011

Sylvester, C. M., Corbetta, M., Raichle, M. E., Rodebaugh, T., Schlaggar, B. L., Sheline, Y. I., Zorumski, C. F., & Lenze, E. J. (2012). Functional network dysfunction in anxiety and anxiety disorders. Trends in Neurosciences, 35(9), 527–535. 10.1016/j.tins.2012.04.012

Taylor, P. J., Gooding, P., Wood, A. M., & Tarrier, N. (2011). The role of defeat and entrapment in depression, anxiety, and suicide. Psychological Bulletin, 137(3), 391–420. 10.1037/a0022935

Thabrew, H., Stasiak, K., Bavin, L.-M., Frampton, C., & Merry, S. (2018). Validation of the Mood and Feelings Questionnaire (MFQ) and Short Mood and Feelings Questionnaire (SMFQ) in New Zealand help-seeking adolescents. International Journal of Methods in Psychiatric Research, 27(3), e1610. 10.1002/mpr.1610

Thomas Yeo, B. T., Krienen, F. M., Sepulcre, J., Sabuncu, M. R., Lashkari, D., Hollinshead, M., Roffman, J. L., Smoller, J. W., Zöllei, L., Polimeni, J. R., Fisch, B., Liu, H., & Buckner, R. L. (2011). The organization of the human cerebral cortex estimated by intrinsic functional connectivity. Journal of Neurophysiology, 106(3), 1125–1165. 10.1152/jn.00338.2011

Van Essen, D. C., Glasser, M. F., Dierker, D. L., Harwell, J., & Coalson, T. (2012). Parcellations and hemispheric asymmetries of human cerebral cortex analyzed on surface-based atlases. Cerebral Cortex (New York, N.Y.□: 1991), 22(10), 2241–2262. 10.1093/CERCOR/BHR291

Varga, T. (2020). Westergaard, D. missCompare: Intuitive Missing Data Imputation Framework.

Williams, L. M., & Goldstein-Piekarski, A. N. (2020). Applying a neural circuit taxonomy in depression and anxiety for personalized psychiatry. In Personalized Psychiatry (pp. 499–519). Elsevier. 10.1016/B978-0-12-813176-3.00042-0

Willinger, D., Häberling, I., Ilioska, I., Berger, G., Walitza, S., & Brem, S. (2024). Weakened effective connectivity between salience network and default mode network during resting state in adolescent depression. Frontiers in Psychiatry, 15. 10.3389/fpsyt.2024.1386984

Wood, A., Kroll, L., Moore, A., & Harrington, R. (1995). Properties of the Mood and Feelings Questionnaire in Adolescent Psychiatric Outpatients: A Research Note. Journal of Child Psychology and Psychiatry, 36(2), 327–334. 10.1111/j.1469-7610.1995.tb01828.x

Xu, J., Van Dam, N. T., Feng, C., Luo, Y., Ai, H., Gu, R., & Xu, P. (2019). Anxious brain networks: A coordinate-based activation likelihood estimation meta-analysis of resting-state functional connectivity studies in anxiety. Neuroscience & Biobehavioral Reviews, 96, 21–30. 10.1016/j.neubiorev.2018.11.005

Zhang, Y., Duan, M., & He, H. (2024). Deficient salience and default mode functional integration in high worry-proneness subject: A connectome-wide association study. Brain Imaging and Behavior, 18(6), 1560–1568. 10.1007/s11682-024-00951-1

Zhang, Z., Zhang, Y., Wang, H., Lei, M., Jiang, Y., Xiong, D., Chen, Y., Zhang, Y., Zhao, G., Wang, Y., Zhang, W., Xu, J., Zhai, Y., An, Q., Li, S., Hao, X., & Liu, F. (2025). Resting-state network alterations in depression: A comprehensive meta-analysis of functional connectivity. Psychological Medicine, 55, e63. 10.1017/S0033291725000303

