## Supplementary Materials for "Shared and unique lifetime stressor characteristics and network connectivity predict adolescent anxiety and depression"

**Supplementary Result 1. Test of normality of LME model residuals before and after square-root transformation**

Besides visually examining the normal Q-Q plots using LME model outputs to assess normality of model residuals, we also conducted two-sample Kolmogorov-Smirnov tests to statistically compare the LME model residuals to the normal distribution. Before square-root transformation, two-sample Kolmogorov-Smirnov tests showed that residuals of the LME models predicting anxiety (*D*s = 0.083, 0.086, 0.099, 0.098, 0.11, 0.099, 0.098, 0.10; *p*s = 0.0038, 0.0027, 0.0023, 0.0028, 0.00086, 0.0026, 0.0027, 0.0023) and depression (*D*s = 0.081, 0.082, 0.099, 0.10, 0.11, 0.10, 0.10, 0.10; *p*s = 0.0052, 0.0050, 0.0023, 0.0011, 0.00096, 0.0016, 0.0014, 0.0016) symptoms did not come from the normal distribution. After square-root transformation, two-sample Kolmogorov-Smirnov tests showed that residuals of the LME models predicting anxiety (*D*s = 0.053, 0.050, 0.053, 0.057, 0.059, 0.055, 0.054, 0.055; *p*s = 0.16, 0.21, 0.29, 0.22, 0.19, 0.25, 0.27, 0.26) and depression (*D*s = 0.034, 0.032, 0.046, 0.047, 0.047, 0.046, 0.044, 0.046; *p*s = 0.68, 0.75, 0.46, 0.44, 0.43, 0.46, 0.51, 0.45) symptoms were not significantly different from the normal distribution.

**Supplementary Table 1**. Bivariate correlations between continuous variables in the LME models

|  | **Anxiety at time 1^a^** | **Anxiety at time 2^b^** | **Depression at baseline** | **Depression at time 1 ^a^** | **Depression at time 2 ^b^** | **Interpersonal loss severity** | **Physical danger severity** | **Entrapment**  **severity** | **Role reversal severity** | **Humiliation severity** | **Mean parental education** | **Baseline Age** | **wFPN FC** | **wDN FC** | **wVAN FC** | **bFPN-DN FC** | **bFPN-VAN FC** | **bDN-VAN FC** |
| --- | --- | --- | --- | --- | --- | --- | --- | --- | --- | --- | --- | --- | --- | --- | --- | --- | --- | --- |
| **Anxiety at baseline** | **0.73****** | **0.64****** | **0.79****** | **0.65****** | **0.61****** | **0.52****** | **0.55****** | **0.73****** | **0.62****** | **0.71****** | -0.10 | **0.18*** | 0.01 | 0.12 | 0.09 | 0.12 | 0.06 | 0.07 |
| **Anxiety at time 1^a^** |  | **0.80****** | **0.68****** | **0.83****** | **0.69****** | **0.40****** | **0.43****** | **0.61****** | **0.52****** | **0.59****** | -0.03 | **0.24***** | -0.02 | 0.11 | 0.08 | 0.11 | 0.04 | 0.07 |
| **Anxiety at time 2^b^** |  |  | **0.58****** | **0.70****** | **0.85****** | **0.32***** | **0.36****** | **0.55****** | **0.44****** | **0.48****** | 0.07 | **0.22**** | -0.04 | 0.03 | 0.03 | 0.05 | 0.02 | 0.02 |
| **Depression at baseline** |  |  |  | **0.69****** | **0.69****** | **0.48****** | **0.51****** | **0.67****** | **0.53****** | **0.65****** | -0.10 | 0.15 | 0 | 0.06 | 0.10 | 0.11 | 0.05 | 0.05 |
| **Depression at time 1^a^** |  |  |  |  | **0.75****** | **0.42****** | **0.44****** | **0.59****** | **0.47****** | **0.56****** | -0.02 | **0.32****** | -0.03 | 0.06 | 0.09 | 0.08 | 0.00 | 0.01 |
| **Depression at time 2^b^** |  |  |  |  |  | **0.33****** | **0.33****** | **0.56****** | **0.42****** | **0.49****** | 0.00 | **0.21*** | -0.06 | -0.01 | 0.04 | 0.02 | 0.01 | 0.01 |
| **Interpersonal loss** |  |  |  |  |  |  | **0.52****** | **0.60****** | **0.67****** | **0.58****** | -0.13 | 0.16 | 0.08 | 0.16 | 0.16 | 0.16 | 0.17 | 0.18 |
| **Physical danger** |  |  |  |  |  |  |  | **0.60****** | **0.63****** | **0.67****** | **-0.17*** | 0.13 | -0.03 | 0.07 | 0.12 | 0.04 | -0.03 | -0.02 |
| **Entrapment** |  |  |  |  |  |  |  |  | **0.80****** | **0.78****** | -0.13 | **0.21*** | -0.02 | 0.11 | 0.11 | 0.10 | 0.04 | 0.09 |
| **Role reversal** |  |  |  |  |  |  |  |  |  | **0.67****** | **-0.16*** | **0.22**** | 0.05 | 0.10 | 0.11 | 0.10 | 0.04 | 0.07 |
| **Humiliation** |  |  |  |  |  |  |  |  |  |  | **-0.21**** | 0.06 | -0.04 | 0.06 | 0.11 | 0.03 | 0.00 | -0.01 |
| **Mean parental education** |  |  |  |  |  |  |  |  |  |  |  | -0.03 | 0.15 | 0.09 | 0 | 0.03 | 0.07 | 0.05 |
| **Baseline age** |  |  |  |  |  |  |  |  |  |  |  |  | 0.13 | 0.16 | 0.04 | **0.22**** | 0.08 | 0.15 |
| **wFPN FC** |  |  |  |  |  |  |  |  |  |  |  |  |  | **0.72****** | **0.73****** | **0.85****** | **0.86****** | **0.74****** |
| **wDN FC** |  |  |  |  |  |  |  |  |  |  |  |  |  |  | **0.78****** | **0.81****** | **0.70****** | **0.64****** |
| **wVAN FC** |  |  |  |  |  |  |  |  |  |  |  |  |  |  |  | **0.73****** | **0.77****** | **0.61****** |
| **bFPN-DN FC** |  |  |  |  |  |  |  |  |  |  |  |  |  |  |  |  | **0.83****** | **0.86****** |
| **bFPN-VAN FC** |  |  |  |  |  |  |  |  |  |  |  |  |  |  |  |  |  | **0.88****** |

Note: All statistics are Pearson's correlation coefficient r. **p*<.05 ***p*<0.01****p*<0.005 *****p*<0.001 a. Time 1 = at 6-month follow-up assessment. b. Time 2 = at 12-month follow-up assessment. FC = functional connectivity. bFPN-DN = between frontoparietal and default network, bDN-VAN = between default network and ventral attention network. bFPN-VAN = between frontoparietal and ventral attention network. wFPN = within frontoparietal network. wDN = within default network. wVAN = within ventral attention network

**Supplemental Table 2**. Demographic information for included and excluded participants from BANDA dataset release 1.1

|  | Included | Excluded | *T*- or χ^2^-Statistic | *p*-value |
| --- | --- | --- | --- | --- |
| *N* | 150 | 65 | - | - |
| Age, mean ± SD | 15.44 ± 0.85 | 15.45 ± 0.84 | -0.062 | 0.951 |
| Sex, (% female) | 92 (61.33) | 49 (75.38) | 3.37 | 0.067 |
| Diagnostic group (%) | | | | |
| Anxiety | 58 (38.67) | 27 (41.54) | 8.19 | 0.017 |
| Control | 52 (34.67) | 11 (16.92) |  |  |
| Depression | 40 (26.67) | 27 (41.54) |  |  |
| Race/ethnicity (%) | | | | |
| White | 121 (80.67) | 49 (75.38) | 6.63 | 0.25 |
| Hispanic | 11 (7.33) | 5 (7.69) | 0.00 | 1.00 |

Note: Anxiety = having a current diagnosis of at least one anxiety disorder and no depressive disorder based on DSM-5. Depression = having a current diagnosis of at least one depressive disorder based on DSM-5.

**Supplementary Figure 1.** Mean lifetime frequency and severity of each stressor characteristic within each diagnostic group at baseline assessment. Anxiety = having a current diagnosis of at least one anxiety disorder and no depressive disorder; Depression = having a current diagnosis of at least one depressive disorder; Control = having no current or lifetime diagnosis of any psychiatric disorder.

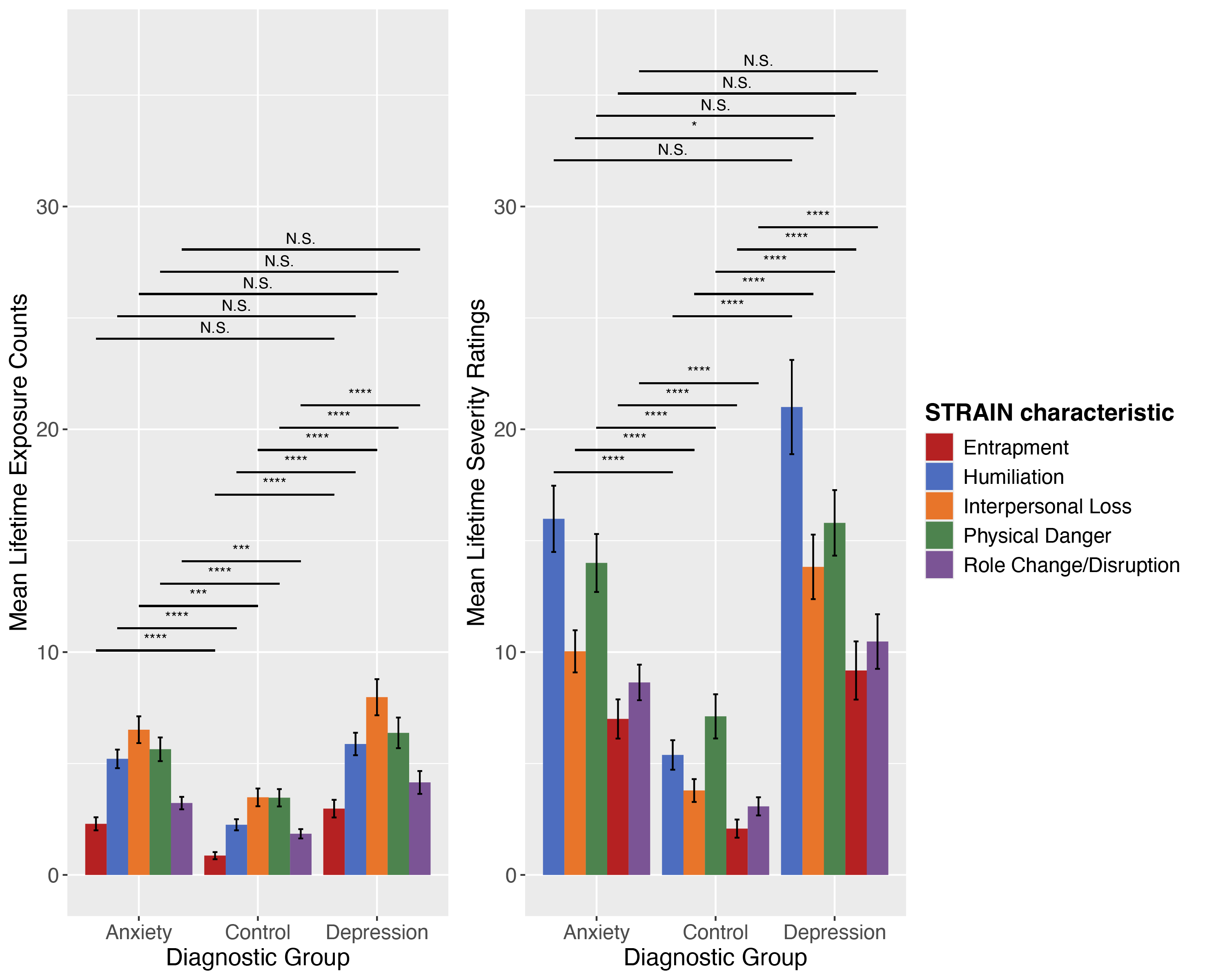

Note: All statistics are based on Tukey’s test. **p*<.05 ***p*<0.01****p*<0.005 *****p*<0.001.

**Supplementary Figure 2**. Mean anxiety and depressive symptoms within each diagnostic group at baseline, 6-month follow-up assessment and 12-month follow-up assessment. Anxiety = having a current diagnosis of at least one anxiety disorder and no depressive disorder; Depression = having a current diagnosis of at least one depressive disorder; Control = having no current or lifetime diagnosis of any psychiatric disorder. Shades around each line represent standard error.

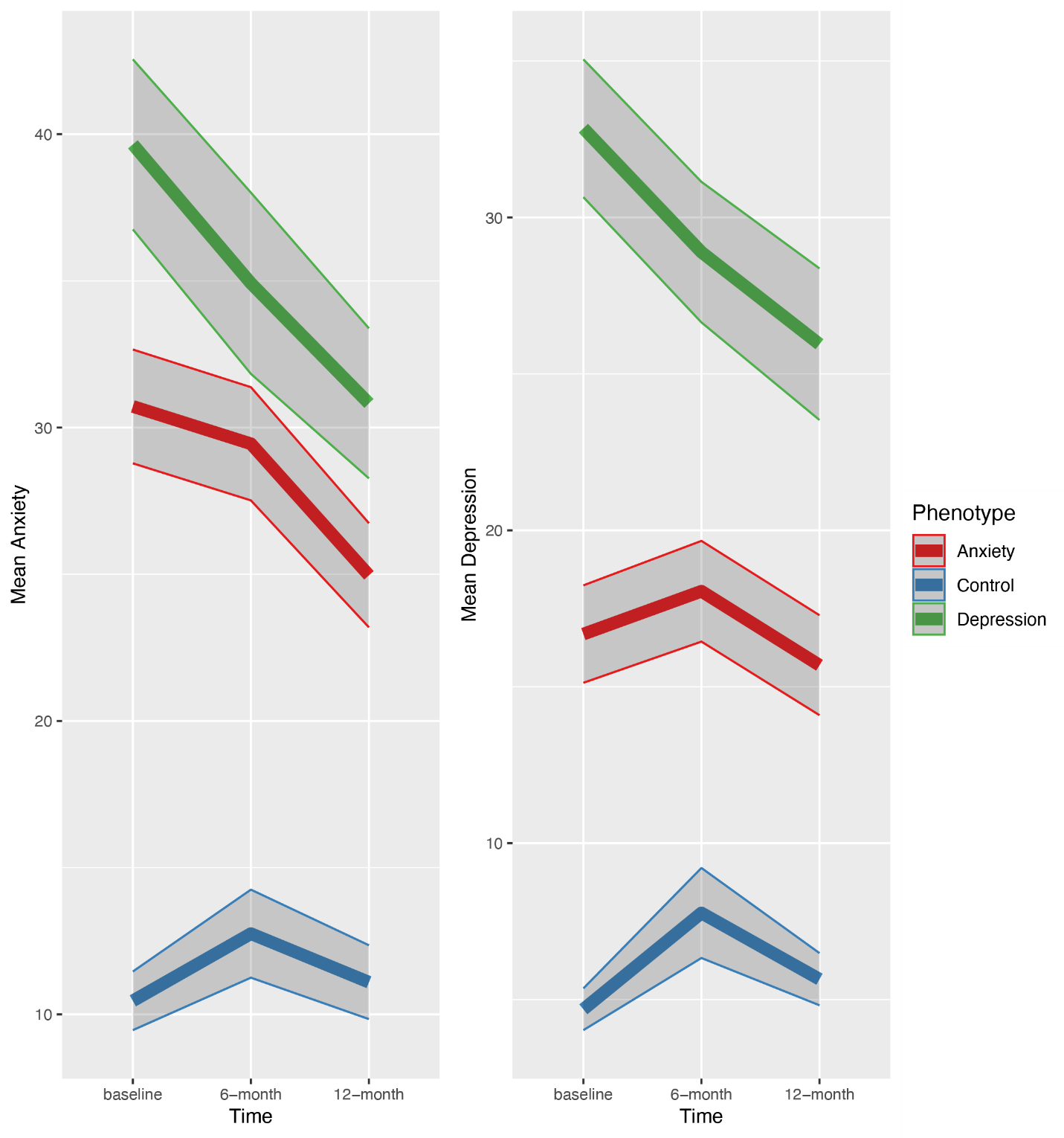

Depression symptoms was computed by the total Mood and Feelings Questionnaire (MFQ), with scores ranging from 0 to 66. Anxiety symptoms were computed by summing the four anxiety subscales (Separation Anxiety Disorder, Social Phobia, Generalized Anxiety and Panic Disorder) from the Revised Children’s Anxiety and Depression Scale (RCADS), ranging from 0 to 93.

**Supplementary Figure 3**. Imputation accuracy (root-mean-square error, mean absolute error, Kolmogorov-Smirnov test statistic) computed by comparing original and imputed data.

Note: Lower values of root-mean-square error, mean absolute error, Kolmogorov-Smirnov test statistic indicate better fit between the original data and imputed data. Some methods (mean imputation, median imputation, missMDA, pcaMethods methods and AmeliaII) do not allow for imputing missing factor variables and therefore inapplicable to our dataset.

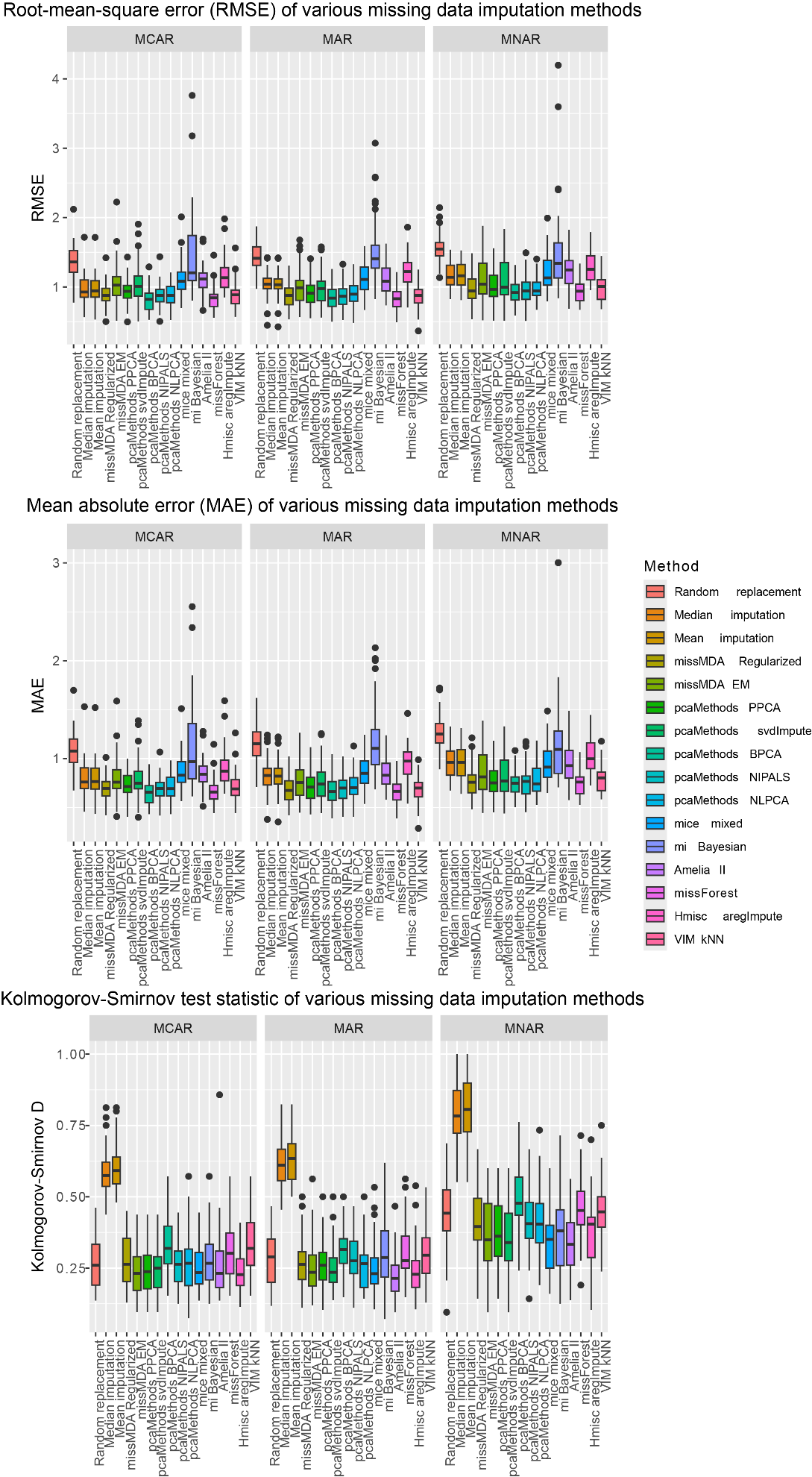

**Supplementary Figure 4**. Bootstrapped correlation coefficients (in red) between the original data and the imputed data. The blue line is a reference line with intercept 0 and slope 1.

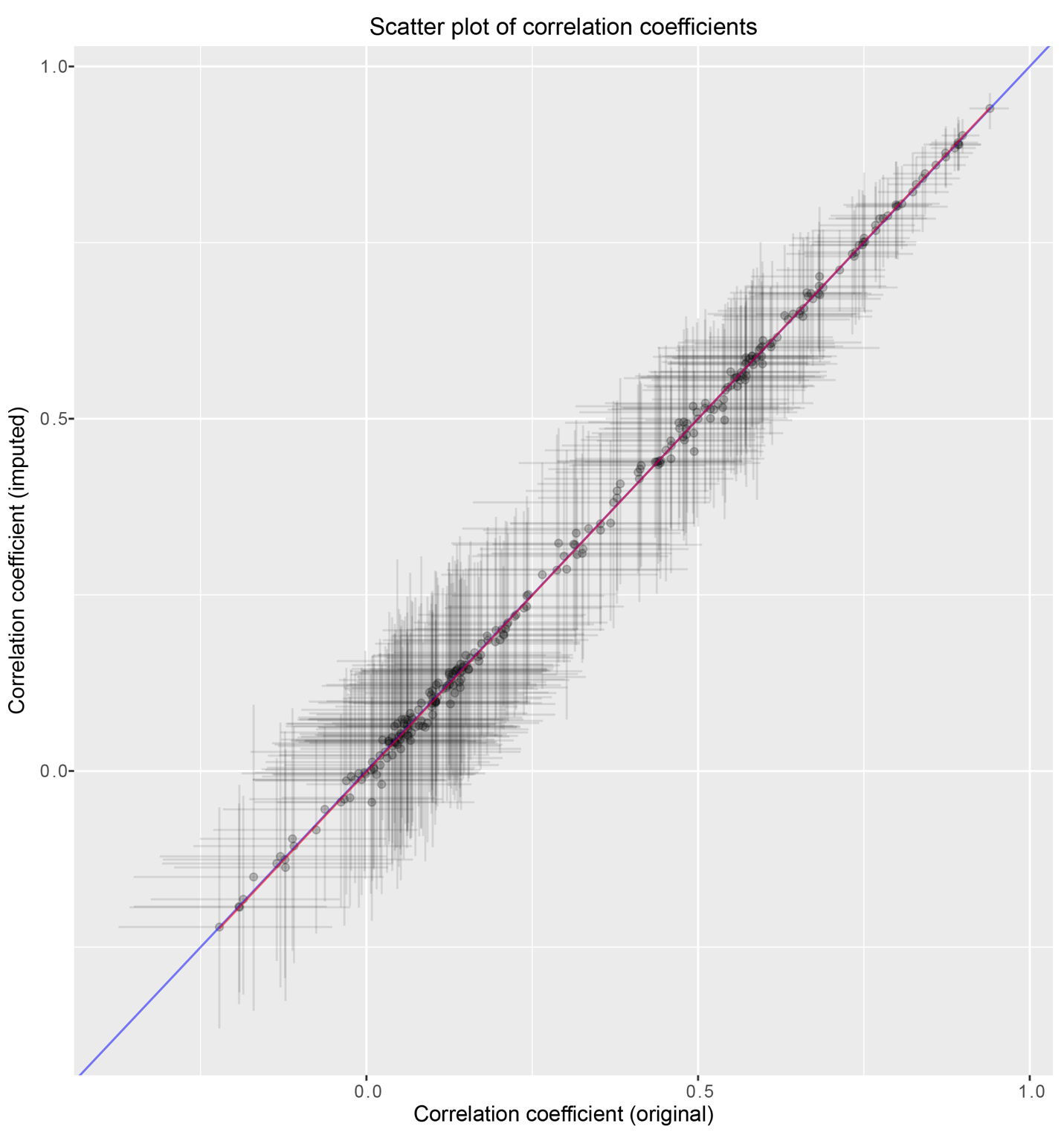

**Supplementary Figure 5**. Q-Q plots for LME models predicting anxiety symptoms before square-root transformation

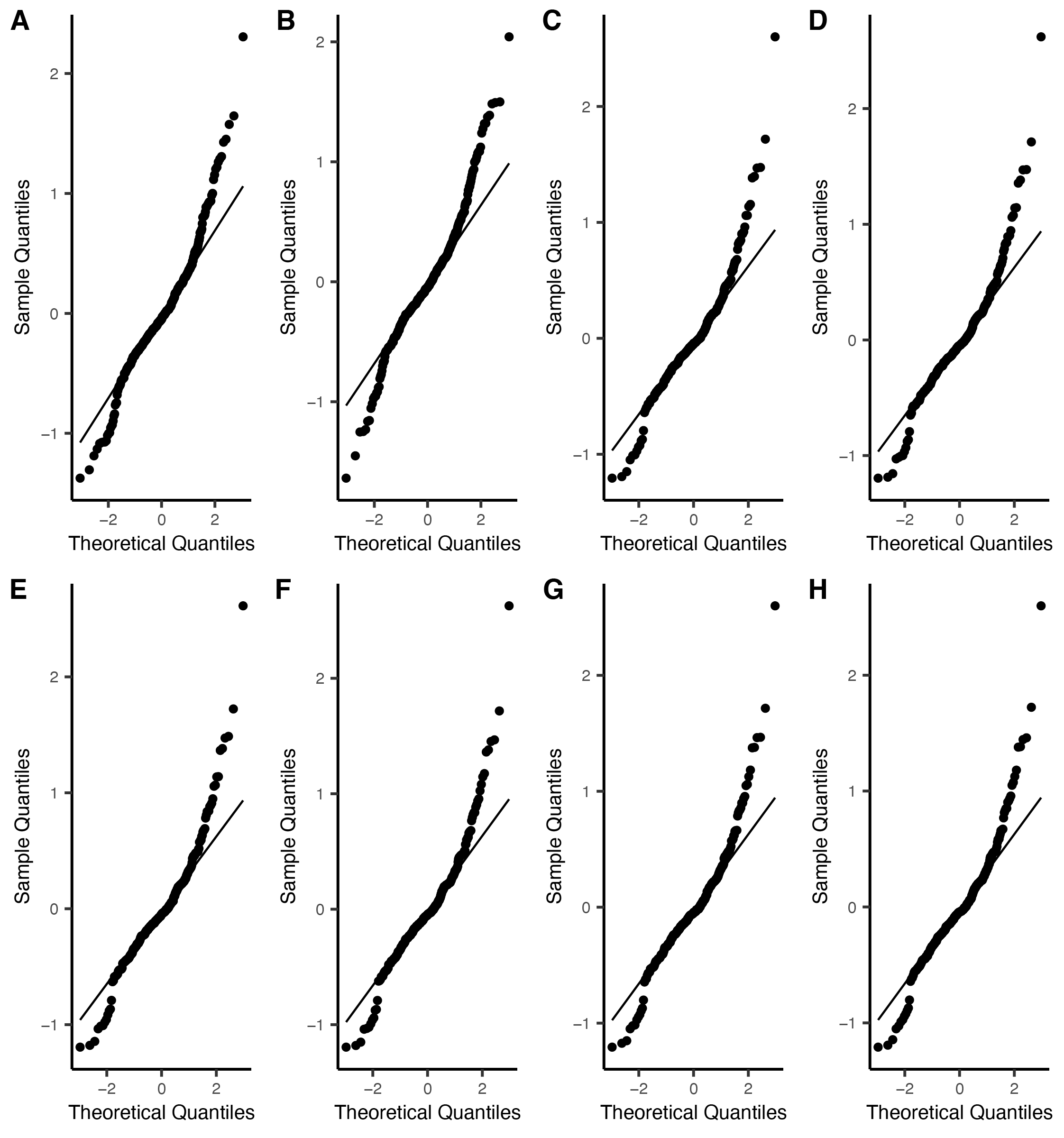

**Supplementary Figure 6.** Heteroscedasticity plots for LME models predicting anxiety symptoms before square-root transformation

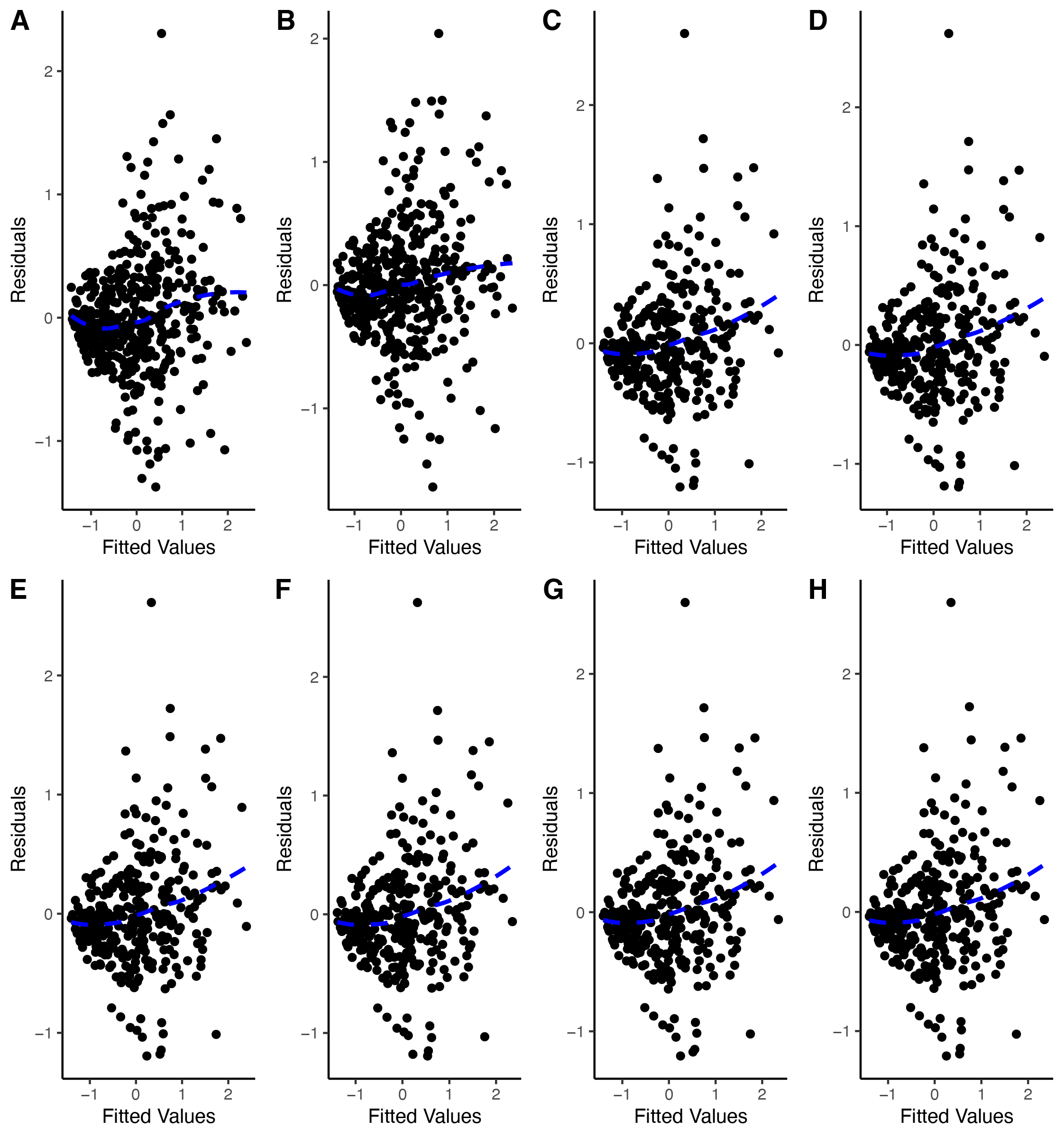

**Supplementary Figure 7**. Q-Q plots for LME models predicting depression symptoms before square-root transformation

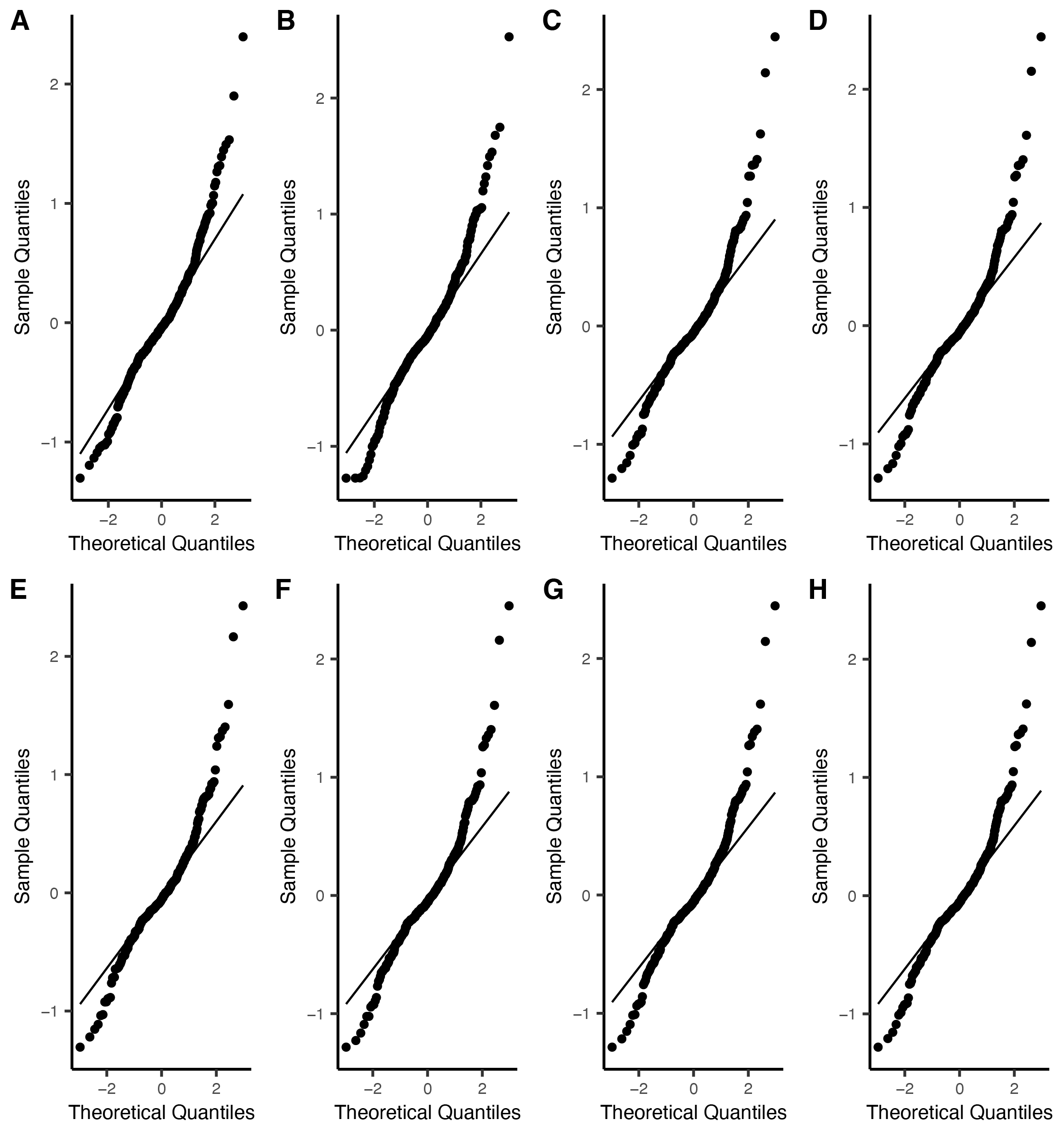

**Supplementary Figure 8**. Heteroscedasticity plots for LME models predicting depression symptoms before square-root transformation

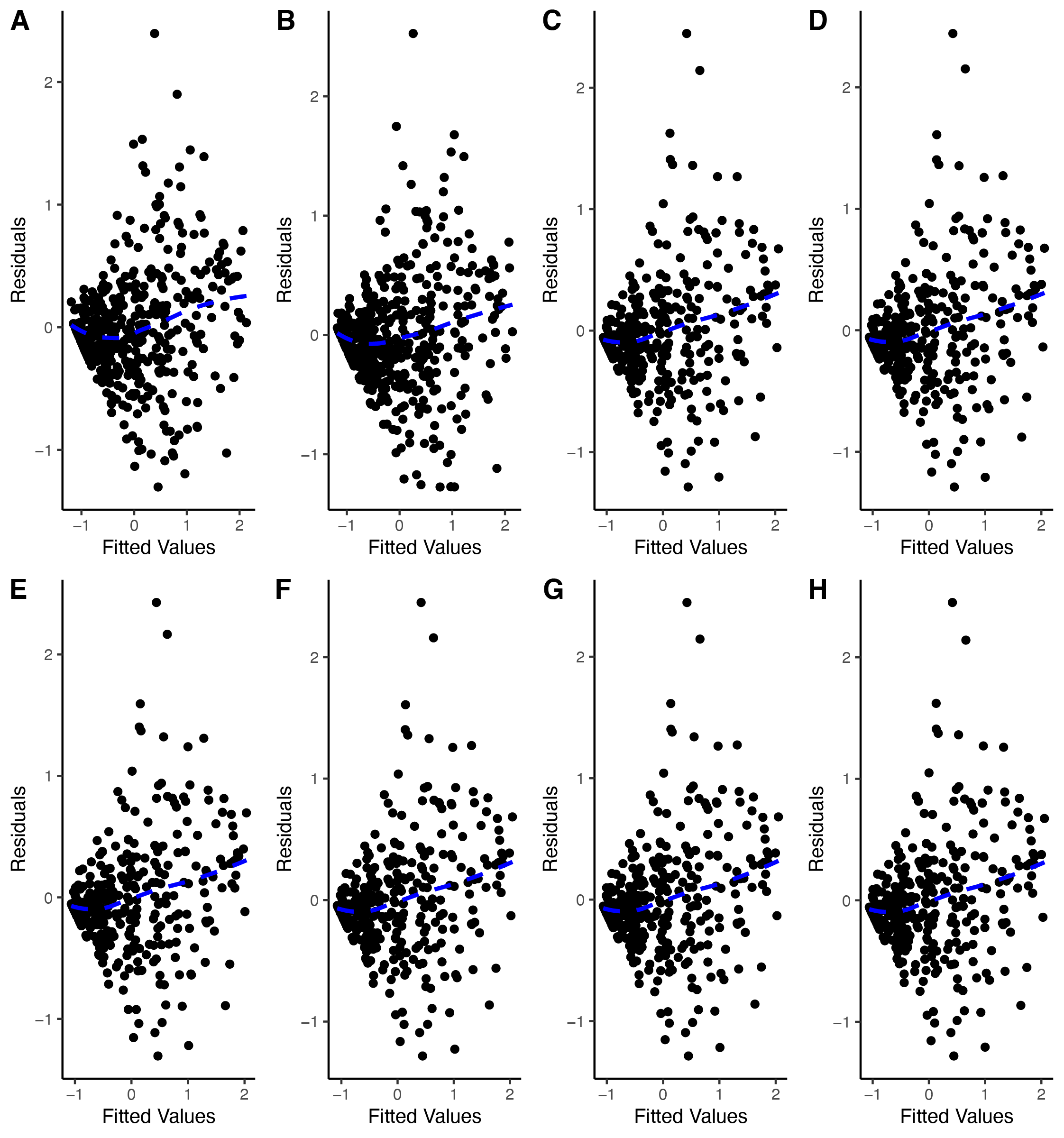

**Supplemental Figure 9**. Q-Q plots for LME models predicting anxiety symptoms after square-root transformation

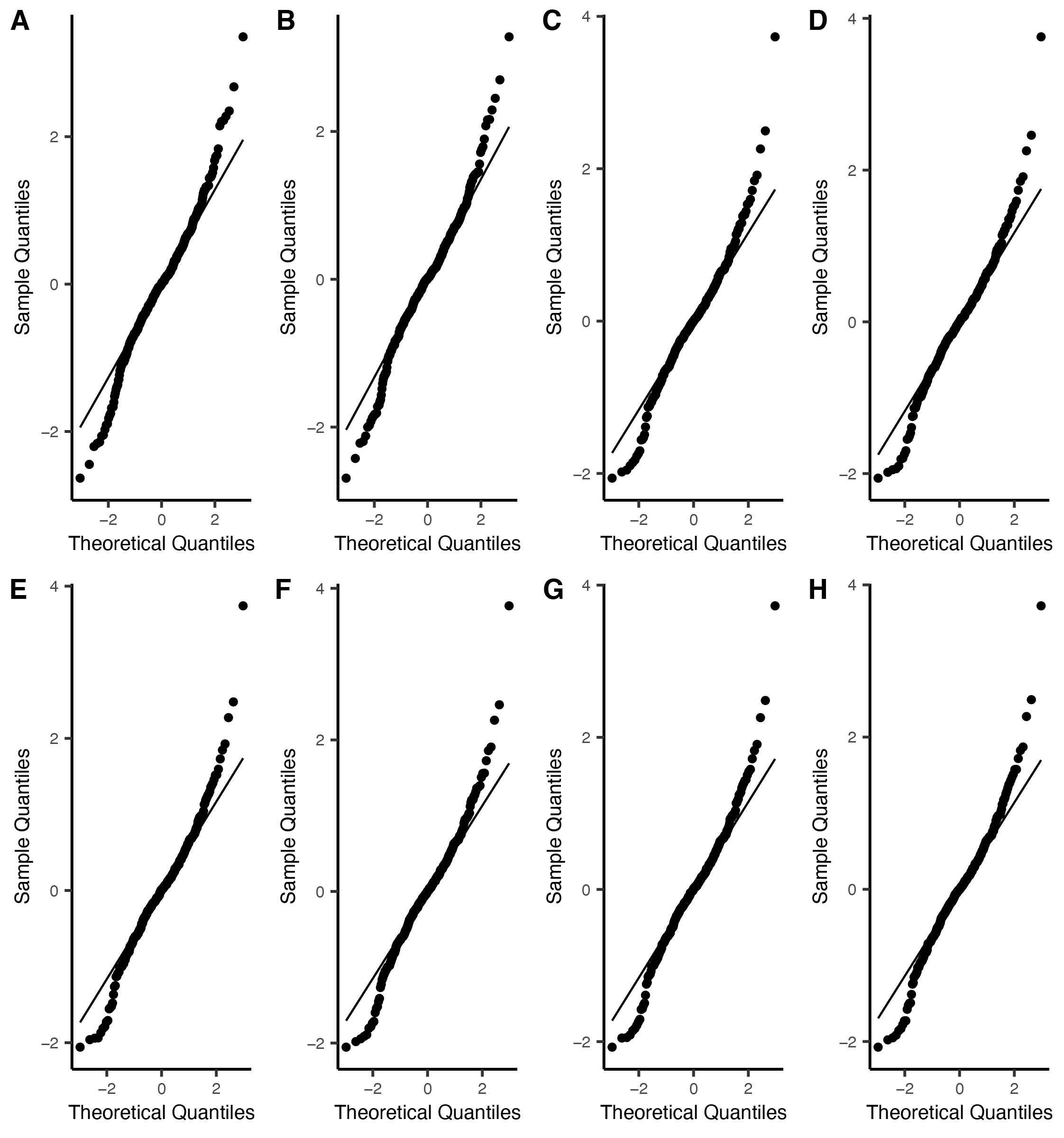

**Supplemental Figure 10**. Heteroscedasticity plots for LME models predicting anxiety symptoms after square-root transformation

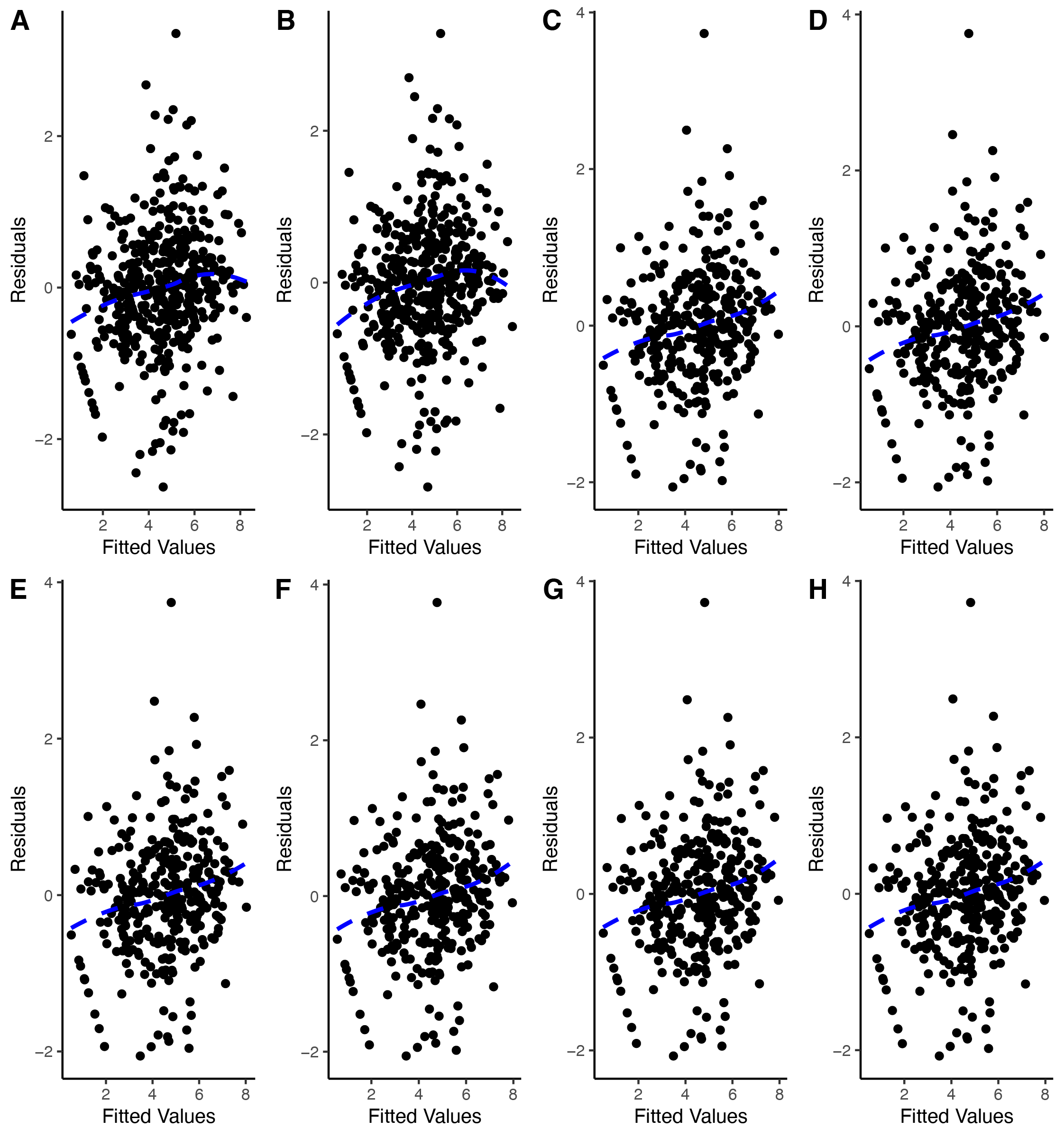

**Supplemental Figure 11**. Q-Q plots for LME models predicting depression symptoms after square-root transformation

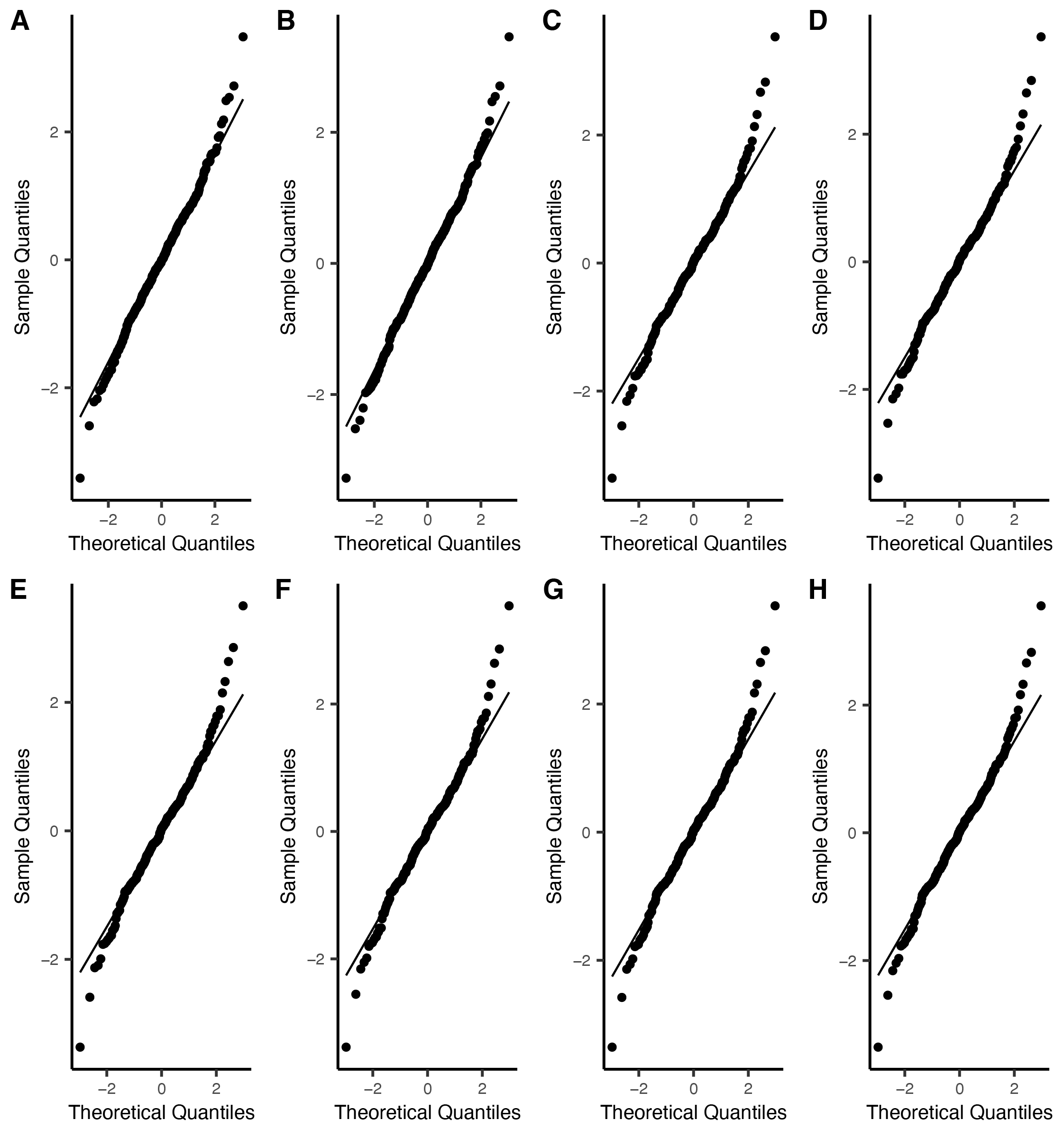

**Supplemental Figure 12**. Heteroscedasticity plots for LME models predicting anxiety symptoms after square-root transformation

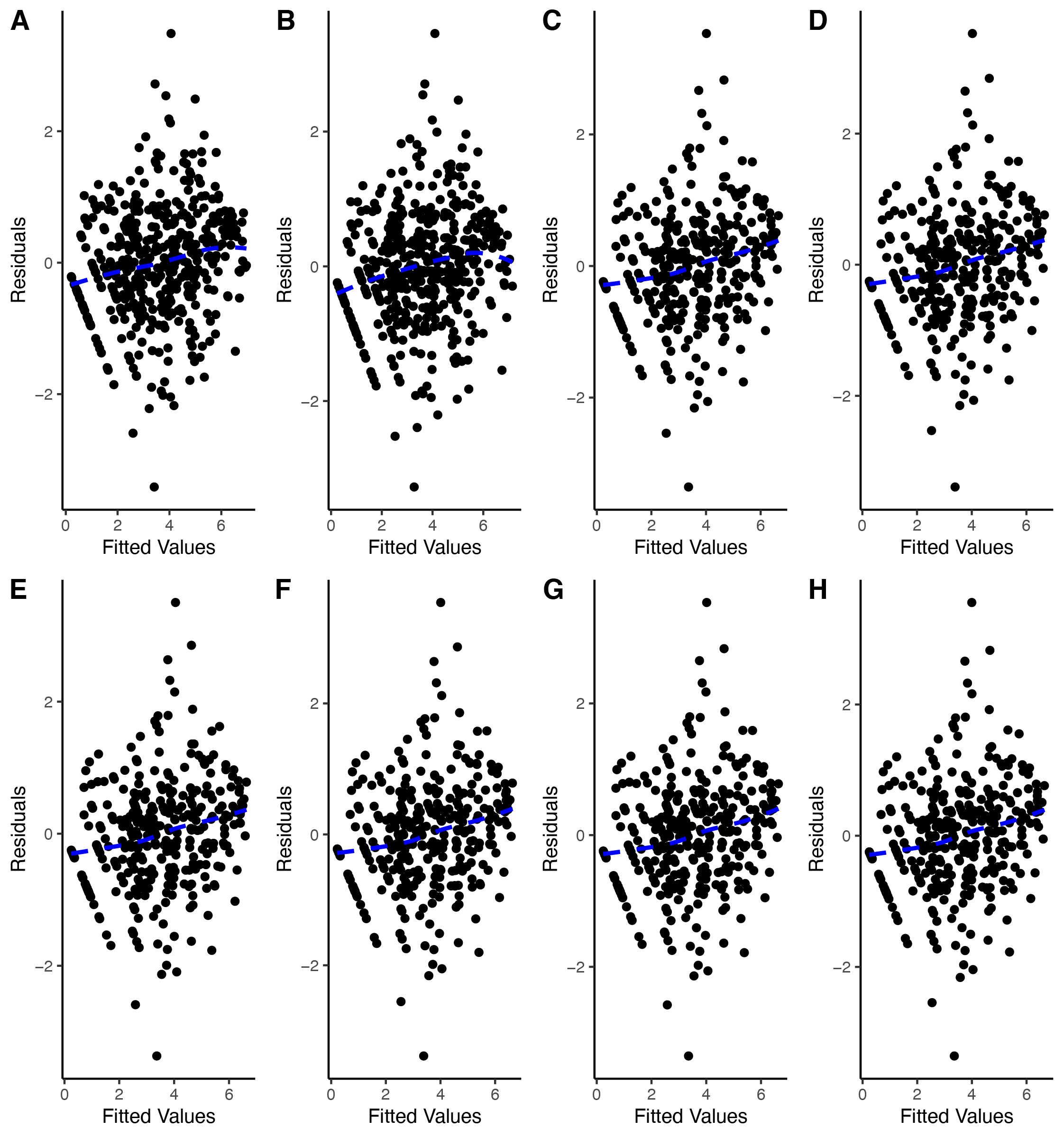

**Supplemental Figure 13.** Comparison of standardized *β* estimates (± 95% confidence intervals) from LME models predicting anxiety and depression symptoms at three time points from total lifetime frequency of five stressor characteristics above and beyond the other symptom at baseline assessment. The vertical dashed line represents *β* = 0. All LME models include a random intercept for each participant and adjust for potentially confounding covariates. Positive *β* estimates indicate that greater stressor frequency is associated with higher symptom scores.

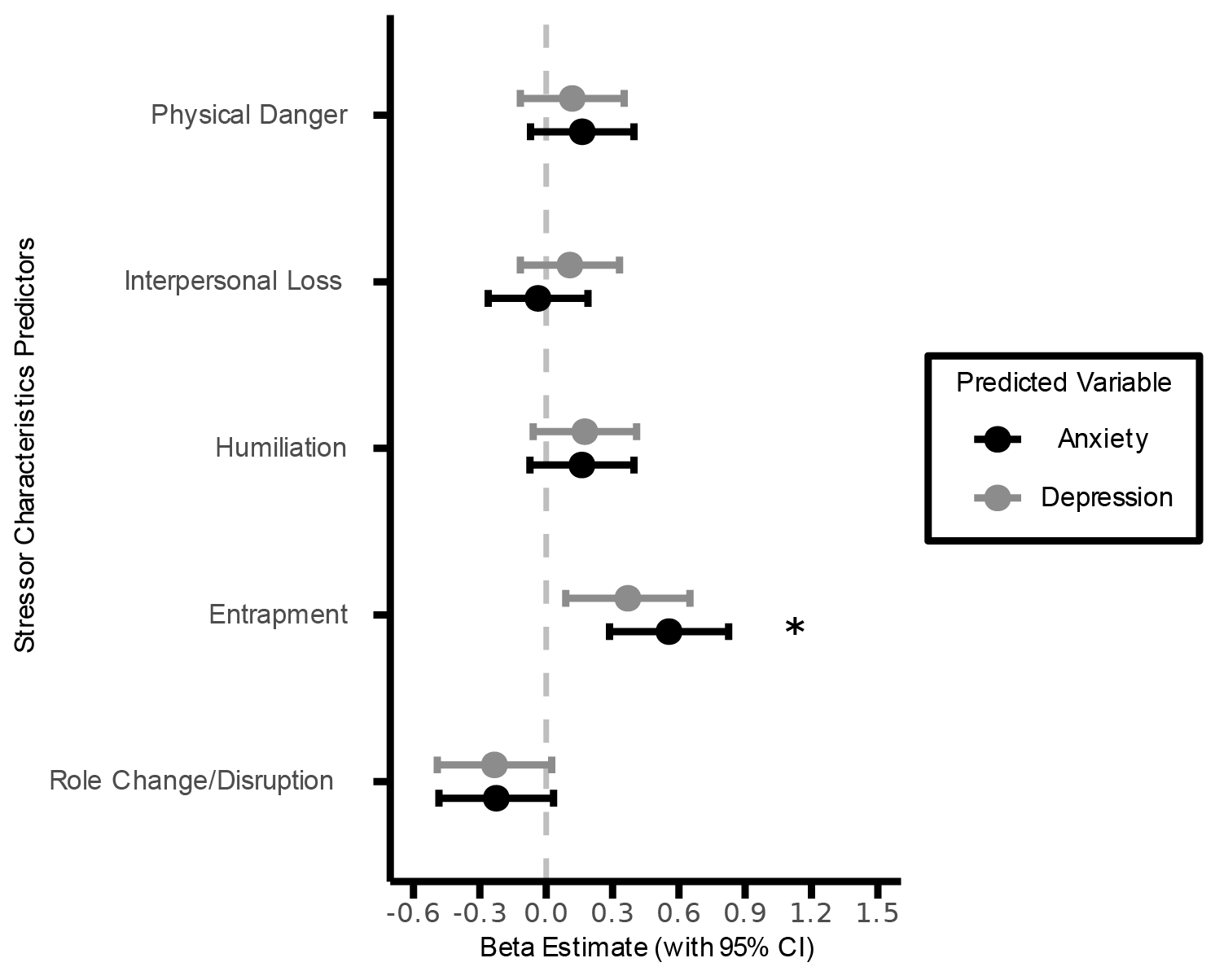

**Supplemental Figure 14**. Comparison of standardized *β* estimates (± 95% confidence intervals) from LME models predicting anxiety and depression symptoms at three time points from total lifetime severity of five stressor characteristics above and beyond the other symptom at baseline assessment. The vertical dashed line represents *β* = 0. All LME models include a random intercept for each participant and adjust for potentially confounding covariates. Positive *β* estimates indicate that greater stressor severity is associated with higher symptom scores. *: q_FDR_ < 0.05

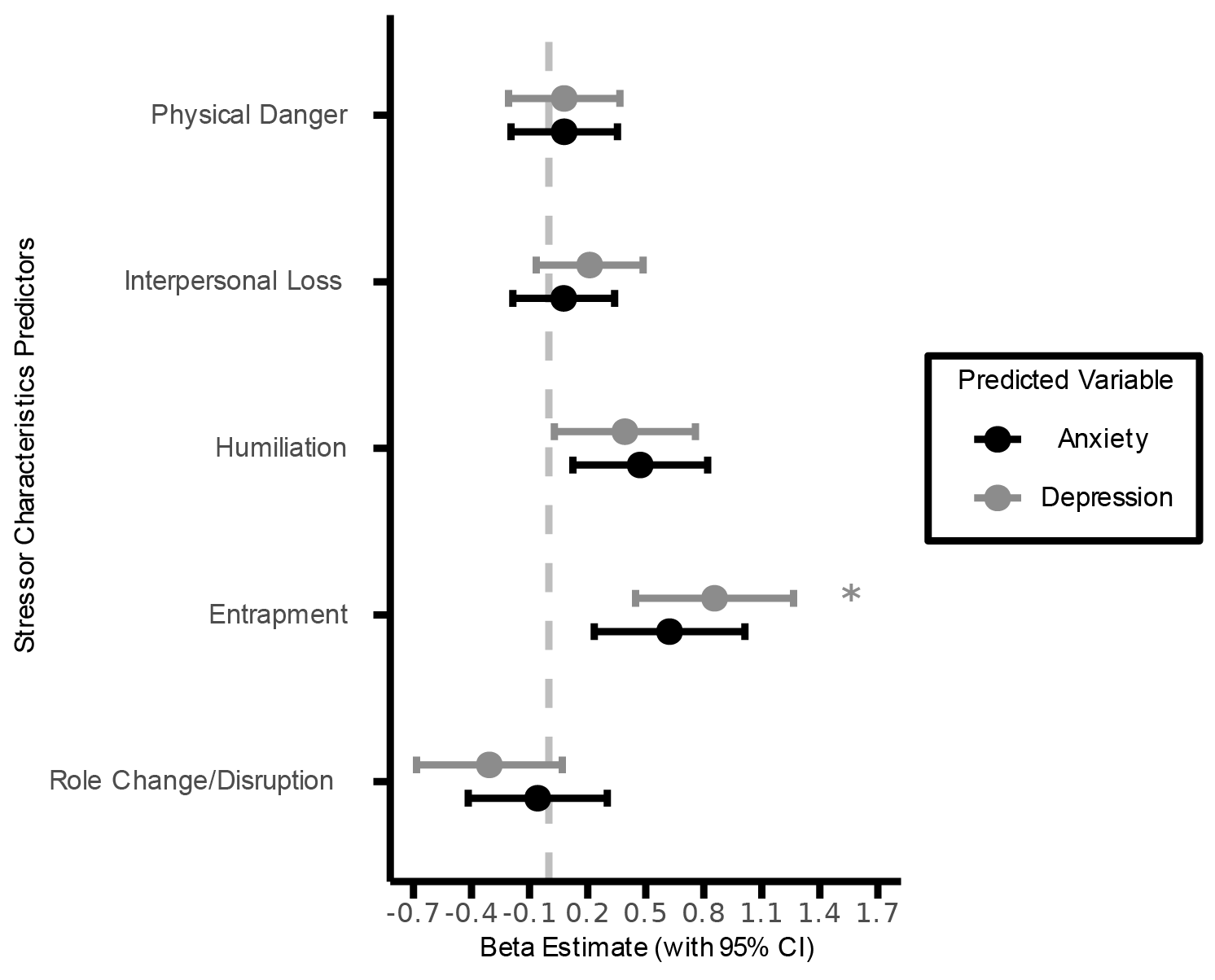

**Supplemental Figure 15**. Comparison of standardized *β* estimates (± 95% confidence intervals) from LME models predicting anxiety and depression symptoms at three time points from RSFC above and beyond the other symptom at baseline assessment. Six RSFC metrics are listed on the y-axis: within-network connectivity of the frontoparietal (FPN), default (DN), and ventral attention (VAN) networks, and between-network connectivity of FPN–DN, FPN–VAN, and DN–VAN. The vertical dashed line represents *β* = 0. All LME models include a random intercept for each participant and adjust for potentially confounding covariates. Positive *β* estimates indicate that greater RSFC is associated with higher symptom scores. *: q_FDR_ < 0.05

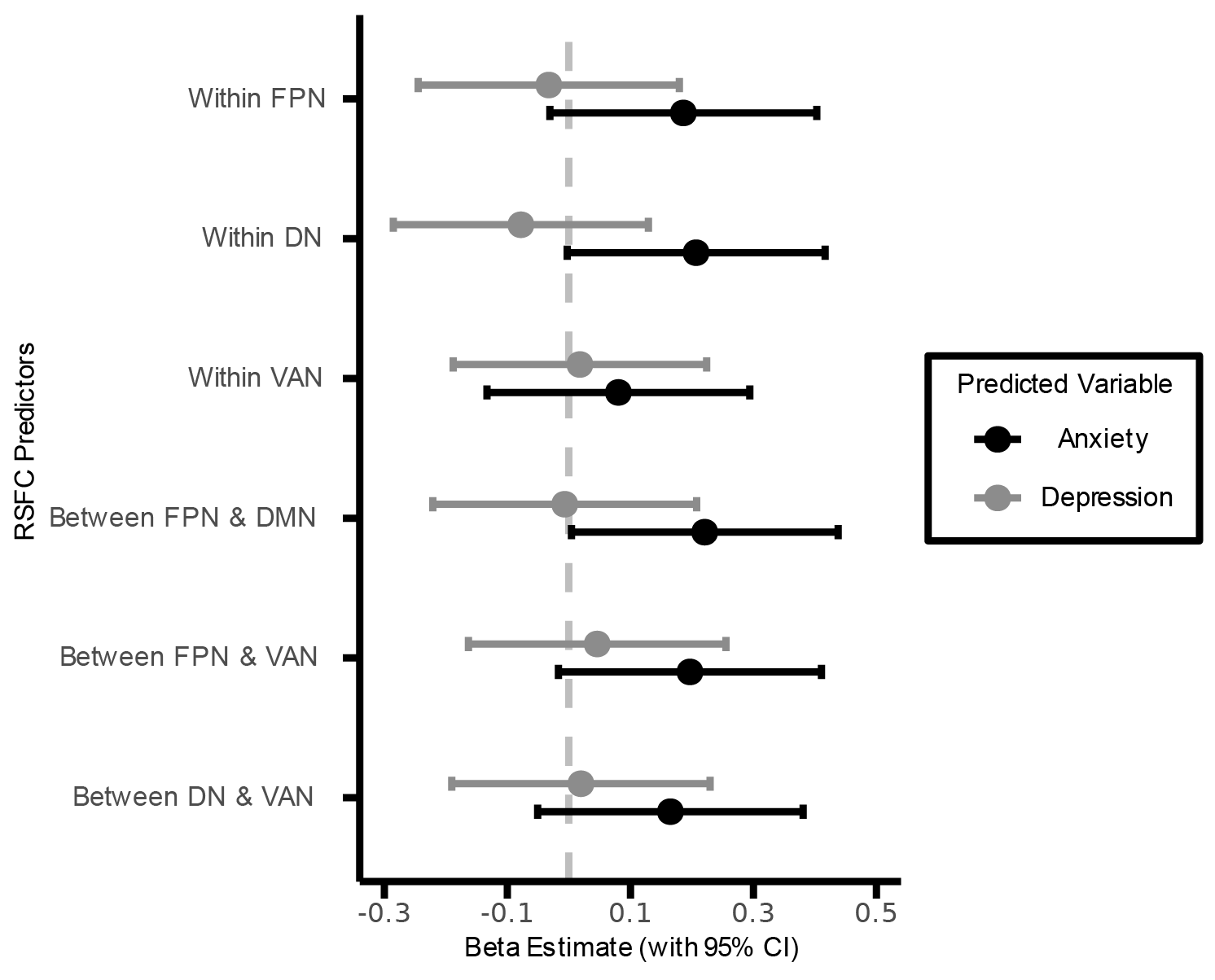
